# A Bidirectional Brain-Fat Body Axis for Pathogen Avoidance

**DOI:** 10.1101/2024.11.02.621634

**Authors:** Yujie Wang, Kokoro Saito, Hiromu Tanimoto, Ilona C. Grunwald Kadow

**Affiliations:** University of Bonn, Institute of Physiology, Faculty of Medicine, Bonn, Germany; Graduate School of Life Sciences, Tohoku University, Katahira 2-1-1, 980-8577 Sendai, Japan

## Abstract

Ingesting pathogens through spoiled food can cause serious harm, including infections, tissue damage, and even death. To prevent these outcomes, many animals have evolved behaviors to avoid consuming harmful pathogens. While pathogen avoidance behavior is conserved across species, the mechanisms linking immune responses of the body with neuron-controlled behavior remain unclear. Building on our previous findings, we here present a new bidirectional body-brain communication between the fat body and the nervous system that drives immune receptor-induced avoidance behavior. We show that immune receptor signaling and a specific antimicrobial peptide (AMP) are essential in both octopaminergic neuromodulatory neurons and the fat body for rapidly reducing pathogen intake after initial ingestion. Mechanistically, the octopaminergic neurons innervate the fly’s fat body where they trigger a calcium response through a specific octopamine receptor. This octopaminergic signal prompts the fat body to release dopamine. In turn, Dop1R1 signaling in output neurons of the mushroom body, the insect higher brain center, drives pathogen avoidance. Together, our data suggest that ingested pathogens are detected by immune receptors in neurons, which, through synaptic connections, trigger the release of dopamine and AMPs from fat cells. While AMPs combat the pathogens, dopamine reduces further ingestion by inducing behavioral changes. This mechanism demonstrates efficient communication between the body and brain, coordinating survival behaviors through systemic dopamine signaling from the fat body to the brain.

## Introduction

Animals, including humans, thrive in a dynamic environment by adapting behavior. Adverse and beneficial elements in the ecological niche therefore have strong effects on fundamental behaviors, such as foraging and feeding, and these effects are surprisingly similar in humans and non-human animals ^1^. Foraging and feeding are complex behaviors that entail many decision processes and are influenced by prior experience. For instance, experiencing the adverse consequences of ingesting toxic or contaminated food will later influence how an animal subsequently forages and feeds ^2^.

Pathogenic microorganisms in spoiled food, just like beneficial microorganisms from the microbiome, influence metabolism and health. They also affect behaviors such as general locomotor activity, sleep and memory ^3^. The mechanisms, signals or neuronal pathways mediating such beneficial or detrimental effects, however, remain to be fully elucidated. Interestingly, the innate immune system emerged as a central player in these processes ^4^. Bacterial peptidoglycans (PGNs) are being recognized by peptidoglycan-recognizing proteins (PGRPs) in various tissues including the nervous system, thereby providing a means for bacteria, good and bad, to influence nervous system function. Notably, these molecular mechanisms appear conserved across a large number of species including *Drosophila melanogaster*, paving the way for genetic approaches aimed at understanding the interaction between immune system, digestive organs and the nervous system ^5,6^.

The PGRP signaling pathway controls the production of various effector peptides and proteins in response to immune challenges. Among these are several key antimicrobial peptides (AMPs), including Diptericin (Dipt), Attacin (Att), Drosocin (Dro), Cecropin, and Defensin. Flies deficient for multiple AMPs are highly susceptible to infections by a wide range of Gram-negative bacteria. AMPs were considered as a general defense mechanism with each individual peptide type contributing in a minor and somewhat redundant manner to the immune response. However, recent work by Hanson et al. (2019) revealed that individual AMPs exhibit a surprisingly high degree of specificity in defending against particular microbes ^7^.

An important source of AMPs in insects is the fat body ^8^. By releasing AMPs into the hemolymph through the activation of immune genes, such as the Toll and immune deficiency (Imd) pathways in response to infections, the fat body helps to combat pathogens like bacteria, fungi, and viruses ^9^. However, the fat body is a multifunctional organ, critical to ensure both metabolic functions and immune defense ^10,11^. Structurally similar to the liver and adipose tissue in mammals, it is distributed throughout the insect’s body, where it surrounds internal organs as well as the nervous system ^8,12^. Metabolically, the fat body serves as a major storage site for lipids, proteins, and carbohydrates, releasing these nutrients during periods of high energy demand such as flight, molting or reproduction ^13^. It plays a central role in regulating diverse metabolic processes, including the synthesis and breakdown of glycogen, lipids, and proteins and produces enzymes essential for maintaining metabolic balance ^8^. Additionally, the fat body functions as a nutrient sensor, coordinating growth, development, and reproduction by modulating hormonal responses ^14^. Notably, recent evidence indicates that fat tissue in mammals, including humans, and flies influences brain function. RNA sequencing, metabolomics as well as genetic experiments in flies posited the putative existence of a fat body-to-brain axis ^15^. However, the mechanistic nature of this axis remains unknown.

Ingestion of pathogens via food can activate such immune responses and the production of AMPs ^16^. Once inside, pathogens can trigger malaise, infection and even damage internal organs ^17^. The onset of these negative effects occurs at least several minutes after food consumption. This timing suggests that animals can adapt their behavior by associating the adverse post-ingestion effects with the food’s sensory characteristics. Indeed, the nematode *C. elegans* can associate an odor with the consumption of pathogenic bacteria and will avoid this odor in the future ^18,19^. In addition, honeybees were shown to associate the post-ingestion effect of a toxin with an odor and avoid this odor in subsequent trials ^20^. Moreover, data from fly larvae show that a food source contaminated with the pathogenic bacterium *Pseudomonas entomophila* (*Pe*) induces lasting avoidance ^21^. A similar earlier study in adult flies showed odor-dependent avoidance of *Pe*-containing food upon prior exposure ^22^. In rodents the commonly used ‘conditioned taste aversion’ assay relies on the lasting association of sweet taste with the delayed onset of chemically or radiation-induced nausea ^23,24^. By now, multiple studies have implicated a specific brain region in the association of negative post-ingestion effects with sensory characteristics of food, including the mushroom body in insects and the amygdala in mammals ^25–30^. Nonetheless, how such negative effects on the body are detected by higher brain centers modulating behavior remains poorly understood ^31^.

Interestingly, previous work in the fly has implicated innate immune signaling in the regulation of neuronal plasticity and behavior. For instance, innate immune receptor PGRP-LC signaling is required for synaptic plasticity at the neuromuscular junction ^32^. Furthermore, the Imd pathway also acts in octopaminergic neurons (OANs) - functionally related to vertebrate noradrenergic neurons- to reduce egg laying upon septic injury induced by PGN or *Escherichia coli* ^33,34^. Remarkably, AMPs have been found to be produced upon associative olfactory long-term memory formation ^35^. Indeed, removal of DiptB from the head fat body and of another AMP, GNBP-like3, from neurons results in memory deficits. In a prior study, we used two well-studied fly pathogens, *Erwinia carotovora carotovora 15* (*Ecc15*) and *Pseudomonas entomophila (Pe)*, to elucidate the mechanisms underlying the post-ingestive effects of contaminated food on food choice behavior ^30^. Surprisingly, despite the negative effects of the two pathogens on the flies’ health, naïve flies prefer the smell of pathogenic bacteria over the one of harmless strains of the same bacteria species. By contrast, flies infected by pathogenic *Ecc15* or *Pe* significantly reduced their consumption of pathogen-contaminated food, demonstrating that flies indeed adapt their behavior to avoid detrimental food sources ^30^. This ability to acquire a dislike for pathogenic bacteria depends on the mushroom body (MB), the insect higher brain center for adaptive behavior and learning ^30,36^. Specifically, we showed that contaminated food avoidance is mediated by the main cells of the MB, the Kenyon cells (KCs), as well as by specific MB output neurons (MBONs), namely MBON01, 03 and 04. In addition, flies deficient for the obligate olfactory receptor *ORCO* also failed to distinguish between pathogenic and harmless strains, indicating that odors are required for the association of spoiled food with its negative consequences ^30^. Our data further showed that innate immune receptor signaling through PGRP-LC and PGRP-LE in OANs mediate this post-ingestion adaptation of feeding behavior ^30^. Based on these data, we proposed a model wherein pathogen ingestion is detected by immune receptors in OANs that convey this information to the MB, where it lastingly modulates feeding behavior, potentially through mechanisms analogous to short-term olfactory associative learning ^30^. The molecular and cellular connection between the body’s immune response and the brain center enabling adaptation of behavior, the MB, remained unclear.

In the present study, we present a new pathway connecting a body organ and a higher brain center that efficiently induces the post-ingestion avoidance of pathogen-contaminated food. In short, our data indicate that pathogen ingestion is detected by immune receptors in OANs and the fat body, which leads to the production of pathogen-targeting AMPs. Importantly, a functional synaptic connection between the nervous system and adipocytes through OANs and an octopamine receptor in the fat body induces a rise in calcium and the subsequent release of dopamine from the fat body. Dopamine then signals to specific MBONs via the dopamine receptor Dop1R1 and induces persistent pathogen avoidance. We propose that neuronally-induced systemic dopamine signaling from the fat body enables efficient communication between brain and body and allows for a fast and global immune response and simultaneous adaptation of survival-relevant behavior.

## Results

### Specific AMPs mediate pathogen avoidance

In our previous work, we provided evidence that the post-ingestion avoidance of food laced with pathogens requires AMPs ^30^. Which AMPs are necessary or where they are produced remained, however, an open question. Given the large number of AMPs and their specificity ^7^, we sought to identify which AMPs are required for flies to adapt their behavior to avoid feeding on harmful bacteria. To this end, we first confirmed the absence of pathogen feeding avoidance in flies lacking most AMPs (ΔAMPs) ^7^. Specifically, we tested the feeding choice with the pathogen *Pe* by comparing the number of cumulative sips from avirulent *Pe GacA*-laced food (5% sucrose) to sips from detrimental *Pe WT*-laced food by using the 1h flyPAD feeding assay ^37^ (Fig. 1A). Remarkably, flies lacking AMPs showed the opposite feeding behavior as compared to controls and preferred to feed on the pathogenic *Pe WT* over the innocuous mutants (Fig.1B). To test how general this mechanism was for Gram-negative bacteria, we tested the flies’ preference for food laced with the harmless *Ecc15 evf* strain over food containing the harmful *Ecc15 pOM1* strain. We again observed the opposite behavior in mutants as seen in controls (Fig. S1A). Flies deficient for AMPs showed a significant preference for the pathogenic *Ecc15 pOM1* strain over the non-pathogenic *Ecc15 evf* strain (Fig. S1A). These data based on two ecologically relevant *Drosophila* pathogens show that flies deficient for AMPs fail to avoid pathogens and are, instead, attracted to these detrimental bacteria. This mutant behavior reflects the olfactory preference of naïve wildtype flies prior to ingestion ^30^. Given the reminiscent results for both bacterial species, we used *Pe* for all subsequent experiments.

**Figure 1.**
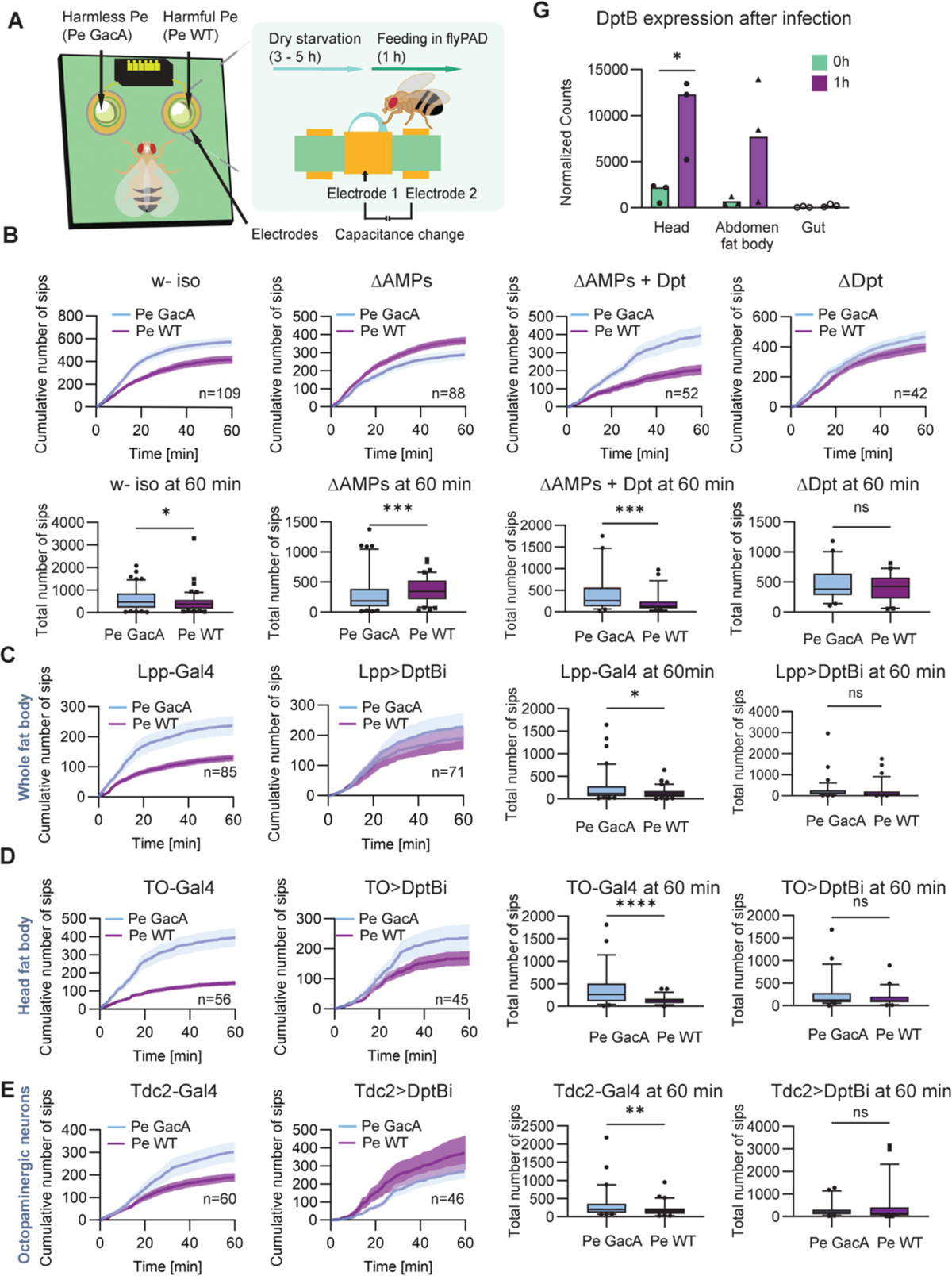
Antimicrobial peptides in fat body and octopaminergic neurons are required for post-ingestion pathogen avoidance. (A) Illustration of the flyPAD feeding choice arena and experimental design. (B) Feeding preferences for food containing harmless and harmful *Pe* bacteria in naïve isogenic w-flies, ΔAMPs flies, flies expressing exclusively Dpt in a ΔAMPs background and Dpt mutant flies, respectively (n (w-) = 109, n (ΔAMPs) = 88, n (ΔAMPs + Dpt) = 52, n (ΔDpt) = 42). Total number of sips at 60 mins; *p-values* calculated via Wilcoxon matched-pairs signed rank test. (C) Feeding preferences for harmless and harmful *Pe* bacteria upon knock-down of DptB in the fat body using Lpp-Gal4;UAS-DptB RNAi (n = 71), and the control flies Lpp-Gal4 (n = 85); experiments were carried out at 28°C to increase RNAi efficacy. Total number of sips at 60 mins; *p-values* calculated via Wilcoxon matched-pairs signed rank test. (D) Feeding preferences for harmless and harmful *Pe* bacteria upon knock-down of DptB in the head fat body using TO-Gal4;UAS-DptB RNAi (n = 45), and the control flies TO-Gal4 (n = 56); experiments at 28°C. Total number of sips at 60 mins; *p-values* calculated via Wilcoxon matched-pairs signed rank test. (E) Feeding preferences for harmless and harmful *Pe* bacteria upon knock-down of DptB in the octopaminergic neurons using Tdc2-Gal4;UAS-DptB RNAi (n = 46), and the control flies Tdc2-Gal4 (n = 60); experiments at 28°C. Total number of sips at 60 mins; *p-values* calculated via Wilcoxon matched-pairs signed rank test. (F) Bar plots depicting normalized counts from DESeq2 for *DptB* expression from head, abdominal fat body and gut after WT flies fed on *Pe WT* for 1 hour. DptB is significantly upregulated in the head upon ingestion of pathogenic *Pe* (*adj. p-value* ≤ 0.1). Three biological replicates were prepared for three tissues (head, fat body and gut) for three independent treatments (exposed to *Pe WT* for 60 mins and control groups). For the groups exposed to *Pe WT,* 75 fly heads, 58 guts and 58 abdominal fat bodies per replicate were used; for the control groups, 45 fly heads, 60 guts and 27 abdominal fat bodies per replicate were used. For all analyses, the statistical notation is as follows: ns (not significant), p > 0.05; * p < 0.05; ** p < 0.01; *** p < 0.001; **** p < 0.0001. Error bars in all panels represent the standard error of the mean (SEM).

Next, we screened flies lacking genes coding for individual or groups of AMPs for their food preference (Fig. S1B-H). We observed that flies lacking DptB, Dro, and Cecropins were unable to distinguish between harmless and pathogenic bacteria, while flies lacking AttA behaved indistinguishable from controls (Fig. 1C, Fig. S1B-H). These data suggest that specific AMPs are involved in pathogen avoidance behavior. We confirmed the function of these AMPs through a rescue experiment where we rescued the two Dpt genes present in the *Drosophila* genome, *DptA* and *-B*, in a *ΔAMPs* background. Flies with this genotype behaved like wildtypes and were able to reduce feeding on harmful forms of *Pe* (Fig. 1C). Interestingly, the ubiquitous (using the driver *actin*-Gal4) downregulation of *DptA* or *DptB* expression by RNA interference gave a contrasted result. Flies deficient for *DptA* retained a wildtype behavior, whereas flies deficient for *DptB* lost the ability to distinguish between avirulent and pathogenic bacteria (Fig. S1B, E). Therefore, we concluded that DiptB is necessary and sufficient for adapting food choice behavior upon pathogen ingestion.

To resolve in which tissue DptB is required for pathogen avoidance behavior, we down-regulated DptB expression in the fat body and in the enterocytes of the midgut since these organs are thought to be at the forefront of the primary immune response ^38,39^. Our data indicated that *DptB* knock-down in the fat body significantly reduces the preference for harmless over harmful bacteria (Fig. 1D, S1I). Moreover, specific down-regulation of *DptB* in the head fat body also led to a strong reduction in preference for harmless *Pe* over its pathogenic form (Fig. 1E), which suggests that the head fat body is a tissue where *DptB* plays a key role in post-ingestion pathogen avoidance. In parallel, we also knocked down *DptB* expression in OANs, as we previously showed that the innate immune receptors, PGRPs, are required in these neurons to mediate this avoidance ^30^. Lack of DptB in OANs also reduced feeding avoidance for pathogenic *Pe WT* (Fig. 1F). However, flies with reduced DptB expression in the enterocytes of the midgut displayed wildtype behavior and still avoided the harmful bacteria (Fig. S1J). We concluded from these experiments that DptB in the fat body and in OANs is essential to mediate post-ingestion pathogen avoidance.

To complement our behavioral assays, we profiled gene expression upon pathogenic bacteria ingestion (1 h after the initial ingestion, Table S1) using RNA-sequencing in different tissues putatively involved in pathogen feeding avoidance: head (including the head fat body), main abdominal fat body and digestive tract. We observed that *DptB* was significantly upregulated in the head, as well as moderately upregulated in the abdominal fat body, albeit with inter-individual variability, but unchanged in the gut, corroborating our behavioral data (Fig. 1G).

Taken together, our data show that a specific AMP, *DptB*, upregulated in the fly head upon pathogen ingestion, is required in both the fat body and in OANs to enable adaptation of feeding behavior upon bacterial infection through contaminated food.

### Octopamine and specific octopamine receptors suppress pathogen feeding

Since PGRP-LC receptor and DptB are required in OANs for post-ingestion pathogen avoidance, we examined the role of OANs, their signaling mode and their targets in enabling the observed behavioral adaptation. We first tested the requirement of octopamine for pathogen avoidance using mutant flies for the gene *tyramine-β-hydroxylase* (*TβH*), the limiting enzyme responsible for converting tyramine into octopamine. In the flyPAD feeding assay, we observed that *TβH* mutant flies failed to avoid pathogenic bacteria and instead fed equally from pathogenic and harmless food sources (Fig. 2A). We concluded that flies relied on octopamine, rather than on tyramine, to suppress feeding on contaminated food.

**Figure 2.**
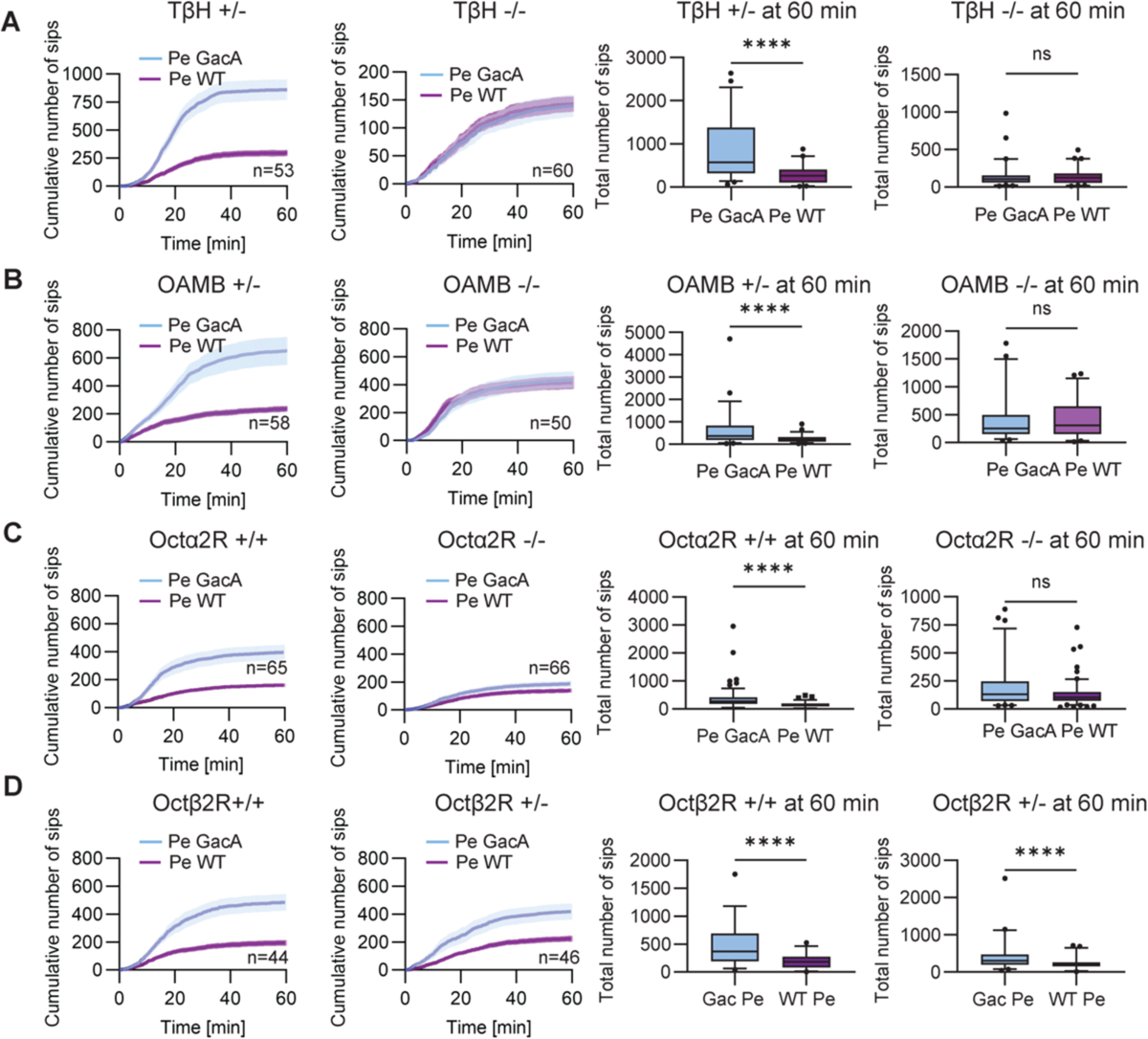
Octopamine and specific octopamine receptors regulate pathogen avoidance behavior. (A) Feeding preferences for harmless and harmful Pe bacteria for TβH mutant flies and heterozygous control flies, n (TβH-/-) = 60, n (TβH+/-) = 53. Cumulative number of sips; mean ± SEM. Total number of sips at 60 mins; *p-values* calculated via Wilcoxon matched-pairs signed rank test. (B) Feeding preferences for harmless and harmful Pe bacteria for OAMB mutant flies and heterozygous control flies: n (OAMB-/-) = 50, n (OAMB+/-) = 58. Cumulative number of sips; mean ± SEM. Total number of sips at 60 mins; *p-values* calculated via Wilcoxon matched-pairs signed rank test. (C) Feeding preferences for harmless and harmful Pe bacteria for Octα2R mutant flies and control flies: n (Octα2R-/-) = 66, n (Octα2R+/+) = 65. Cumulative number of sips; mean ± SEM. Total number of sips at 60 mins; *p-values* calculated via Wilcoxon matched-pairs signed rank test. (D) Feeding preferences for harmless and harmful Pe bacteria for Octβ2R heterozygous mutant flies (Octβ2R-/+) and control flies: n (Octβ2R-/+) = 46, n (Octβ2R+/+) = 44. Octβ2R homozygosity was lethal. Cumulative number of sips; mean ± SEM. Total number of sips at 60 mins*; p-values* calculated via Wilcoxon matched-pairs signed rank test. For all analyses, the statistical notation is as follows: ns (not significant), p > 0.05; * p < 0.05; ** p < 0.01; *** p < 0.001; **** p < 0.0001. Error bars in all panels represent the standard error of the mean (SEM).

To understand how octopamine mediates this avoidance of infected food, we examined the requirement of octopamine receptors. Five octopamine receptors have been previously identified in the fly genome: OAMB, Octα2R, Octβ1R, Octβ2R and Octβ3R ^40^. We initially screened null mutant flies for each of these receptors and found that OAMB and Octα2R were required for pathogen avoidance (Fig. 2B-D, Fig. S2A-B). In addition, we analyzed Octß2R heterozygous mutant flies, as homozygous mutant flies did not survive. We did not observe a significant change in feeding preference in Octß2R heterozygous mutant flies (Fig. 2D).

The strong contribution of OAMB to post-ingestion pathogen avoidance as well as its prior implication in Pavlovian learning and memory prompted us to first investigate its site and mode of action. OAMB is well known for its role in classical olfactory conditioning in dopaminergic neurons innervating the MB (MB-DANs) ^41,42^. In addition, OAMB is required in DANs for appetitive learning ^43–45^. Therefore, we wondered whether it could function as a mediator for pathogen avoidance through the MB network since we previously showed that post-ingestion pathogen avoidance requires the activity of MB intrinsic KCs, as well as of specific MBONs ^30^ Thus, we used RNA interference to knock-down OAMB in both main clusters of MB-DANs, PAM and PPL1 (Fig. S2C-E). As a positive control, we used ubiquitous knock-down of OAMB with the actin-Gal4 driver which confirmed the phenotype observed in *OAMB* mutant flies (Fig. S2C). By contrast, knock-down of OAMB in either PAM or PPL1 neurons did not interfere with the fly’s ability of distinguishing between harmful and harmless bacteria and the flies still avoided the pathogen-containing food source (Fig S2D,E).

Taken together, these results indicate that octopamine and specific octopamine receptors are required for the fly’s ability to suppress feeding on pathogenic bacteria. In addition, they also suggested that octopamine receptor signaling may not be required in MB-DANs.

### An octopaminergic neuron-to-fat body connection is involved in pathogen avoidance

Considering our results above, we tested an alternative theory to explain the observed requirement of octopamine and its receptors. Our results indicated an essential role for the fat body in post-ingestion pathogen avoidance behavior (Fig. 1D, E). Interestingly, prior work uncovered a neuro-adipose junction between sympathetic noradrenergic neurons and adipose tissue in rodents ^46^. Given that octopamine is thought to be the insect functional equivalent of noradrenaline in mammals, we wondered if such an axis could exist in flies and possibly contribute to post-ingestion pathogen avoidance behavior. In line with this hypothesis, results in locusts demonstrated that octopamine stimulates an increase in fat cell cAMP levels as well as the production and release of lipids from the fat body ^47^.

Thus, we next asked if octopamine receptors are indeed expressed in the fat body. To this end, we used a red fluorescent protein to visualize the expression of OAMB, Octβ2R and Octβ3R in the fat body. Specifically, we expressed UAS-myr::mRFP through the Gal4 knock-in reporter OAMB-T2A-Gal4, Octβ2R-T2A-Gal4 or Octβ3R-T2A-Gal4 ^48^ and then imaged the fat body in the head and abdomen using confocal microscopy (Fig. 3A, S3A). Particularly, Octβ2R showed strong mRFP signals in the head fat body (Fig. 3A), which suggested that Octβ2R is expressed in the head fat body. Given the receptor expression in the fat body, we assessed the potential innervation of the fat body by OANs by genetically targeting the expression of two types of fluorescent reporter proteins in both cell types. We observed that OAN neurites ran near the head and abdominal fat body (Fig. 3B, S3B). To analyze whether this spatial proximity is compatible with synaptic connections, we used the GRASP-method (GFP reconstituted across synapse partners) ^49^. Using this method, we could identify reconstituted GFP signals in both head and abdominal fat bodies, at the level of the cell membrane, consistent with synaptic innervation of the fat bodies by OANs (Fig. 3C).

**Figure 3.**
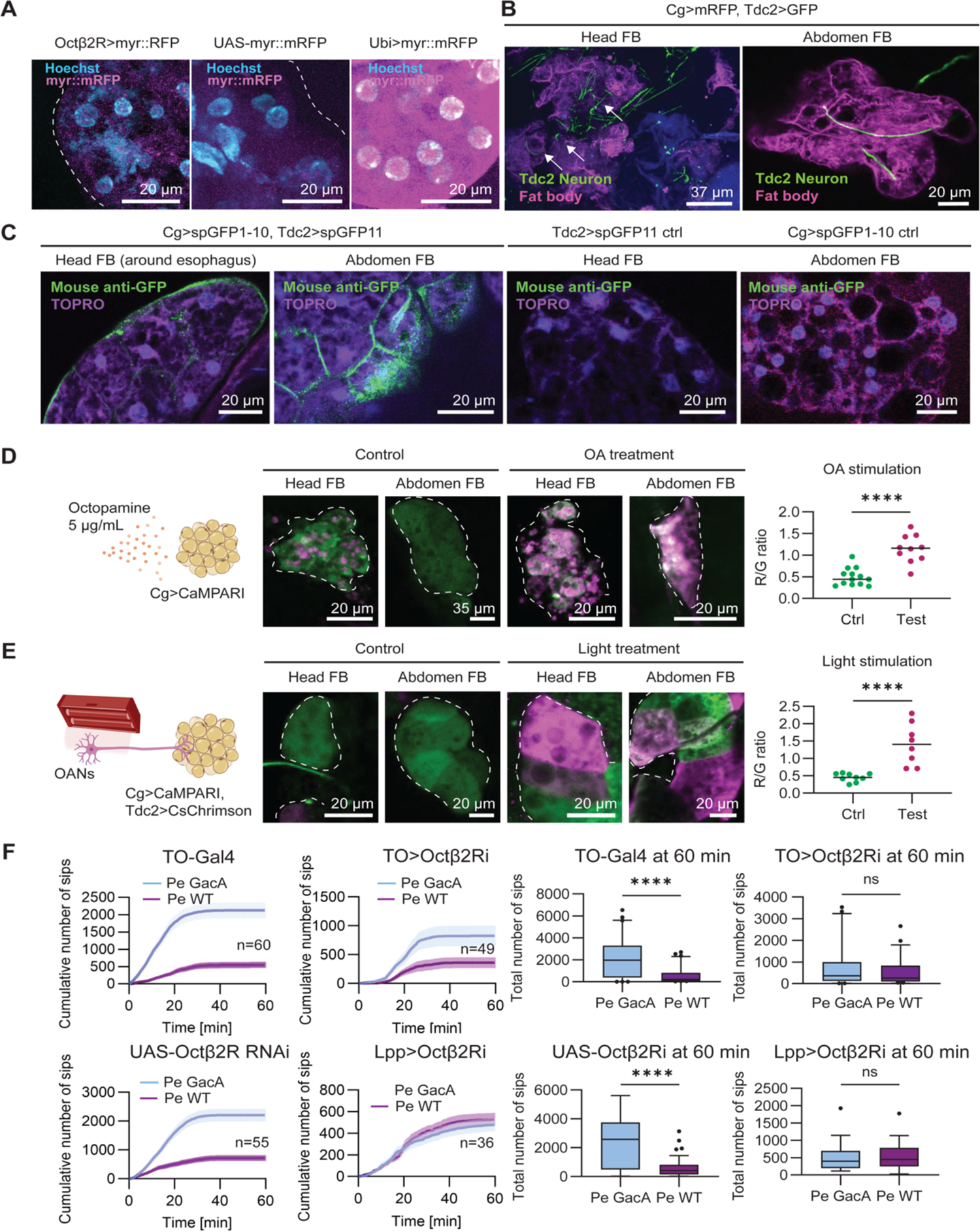
A direct neuronal connection between octopaminergic neurons and the fat body enables post-ingestion pathogen aversion. (A) Representative maximum intensity Z-projection images of Octβ2R expression (Octβ2R-Gal4;UAS-my::mRFP; magenta) in the head fat body, effector control (UAS-myr::mRFP) and positive control (Ubi-Gal4; UAS-myr::mRFP). Nuclei are labelled with Hoechst33342 staining (cyan). Scale bar = 20 μm. (B) Immunostainings showing anatomical proximity of OANs (green) and fat body cells (magenta) in both fly head and abdomen (Cg-Gal4>UAS-mRFP; Tdc2-LexA>LexAop-GFP). Nc82 antibody staining was used as counterstaining for the fly brain as it labels presynaptic active zones (blue). Scale bar = 20 μm. (C) GRASP fluorescence (green) labels the boundary between fat body cells expressing CD4-spGFP1-10 and OANs expressing CD4-spGFP11. Fat body cells were visualized using autofluorescence with 561 nm light excitation (magenta) and fat body nuclei using TOPRO3 staining (purple). Tdc2-Gal4>UAS-spGFP11 and Cg-Gal4>UAS-spGFP1-10 were used as negative controls, respectively. Scale bar = 20 μm. (D) Representative maximum intensity Z-projection images of fat bodies expressing CaMPARI2-L398T after octopamine (OA) bath application (5 µg/mL for 8–10 min) or saline control. Photoconversion was significantly higher in the OA-treated group (n=10; 5 head and 5 abdominal fat bodies from 5 flies) compared to the control (n=13; 5 head and 8 abdominal fat bodies from 8 flies), *p* < 0.0001, Mann-Whitney test. (E) Representative maximum intensity Z-projection images of the fat body expressing CaMPARI2 upon optogenetic stimulation of octopaminergic neurons (OANs) via Cs-Chrimson activation. Photoconversion was significantly higher in the light-treated group (n = 8, from 4 head and 4 abdominal fat body tissues, 4 flies) compared to the control (n = 9, from 4 head and 5 abdominal fat body tissues, 5 flies). p = 0.0159, calculated via Mann-Whitney test. (F) Feeding preferences for harmless and harmful Pe bacteria upon knock-down of Octβ2R in the head fat body and whole fat body: n (TO-Gal4;UAS-Octβ2R RNAi) = 49, n (Lpp-Gal4;UAS-Octβ2R RNAi) = 36 respectively; the control flies, n (TO-Gal4) = 60, n (UAS-Octβ2R RNAi) = 55. Cumulative number of sips; mean ± SEM. Total number of sips at 60 mins; p-values calculated via Wilcoxon matched-pairs signed rank test For all analyses, the statistical notation is as follows: ns (not significant), p > 0.05; * p < 0.05; ** p < 0.01; *** p < 0.001; **** p < 0.0001. Error bars in all panels represent the standard error of the mean (SEM).

However, although GRASP signals indicate proximity between cells and cell membranes, it does not directly demonstrate the existence of functional synapses. We therefore investigated the functionality of these potential connections between OANs and the fat body by using the calcium-modulated photoactivatable ratiometric integrator CaMPARI2-L398T ^50^ to test the response of the fat body to octopamine application and to the activation of OANs. First, we observed a significant increase in CaMPARI UV-light induced photoconversion after 8-10 minutes octopamine (5 µg/mL) bath application onto an explant of the fat body (Fig. 3D). This result indicated an increase in intracellular calcium concentration in fat cells in response to octopamine. Second, we used optogenetics to photoactivate OANs and thereby stimulate the release of octopamine onto the explant fat body (Fig. 3E, S3C). Again, we observed an increase in CaMPARI photoconversion in head and abdominal fat body following OANs photostimulation compared to the no-light stimulation control group (Fig. 3E). These data provide strong support for the presence of functional synaptic connections between OAN and the fat body in the fly.

To test whether these functional connections play a role in post-ingestion pathogen feeding avoidance, we turned to our flyPAD choice assay and removed octopamine receptors from the fatbody using RNAi knock-down. Surprisingly, RNAi-mediated reduction of OAMB in the fatbody caused lethality. Unfortunately, this lethality prevented us from testing its role in behavior. However, when we knocked down the receptor Octβ2R in all fat body tissue or exclusively in the head fat body, the flies survived but no longer developed an aversion to the harmful bacteria (Fig. 3F).

Taken together, our data suggest that, like noradrenergic neurons in mammals, insect octopaminergic neurons form functional synaptic connections with fat tissue. Notably, this innervation of the fly fat body and receptor Octβ2R signaling in fat cells are required for the post-ingestion avoidance of pathogen-contaminated food.

### Dopamine from fat body but not from DANs is required for acquired pathogen aversion

Having shown that a synaptic connection from OANs to the fat body modulates pathogen avoidance post-ingestion, we next sought to unravel the mechanism by which this connection can change behavior. Specifically, we wondered how changes in fat cells can induce a behavioral change.

In addition to an increase in AMP expression in the fat body after oral infection, our analysis of the RNAseq data also revealed an increase in *pale* (*ple*) expression in the fat body. *Ple* encodes tyrosine hydroxylase (TH), the limiting enzyme for dopamine synthesis (Table S1, Fig. S4A). In spite of our result that octopamine receptor OAMB was not required in MB-DANs (see Fig. 2), this finding prompted us to reconsider a putative involvement of dopamine. First, we examined whether MB-DANs are required in flies to distinguish between harmless and harmful forms of *Pe* by optogenetically inhibiting MB-DAN activity during feeding behavior ^49,50^. Consistently with our previous data ^30^, optogenetic inhibition of both PAM and PPL1 DAN clusters did not modify the flies’ post-ingestion avoidance of food infected with pathogenic bacteria (Fig. 4A-B, S4B). To confirm this result with another approach, we down-regulated the expression of *ple* (TH) through RNA interference to prevent with the production of dopamine in these neurons. Once again, there was no difference in behavior between the test and control groups (Fig. S4E, F). We therefore concluded that DANs are not essential for the distinction between harmless and harmful bacteria.

**Figure 4.**
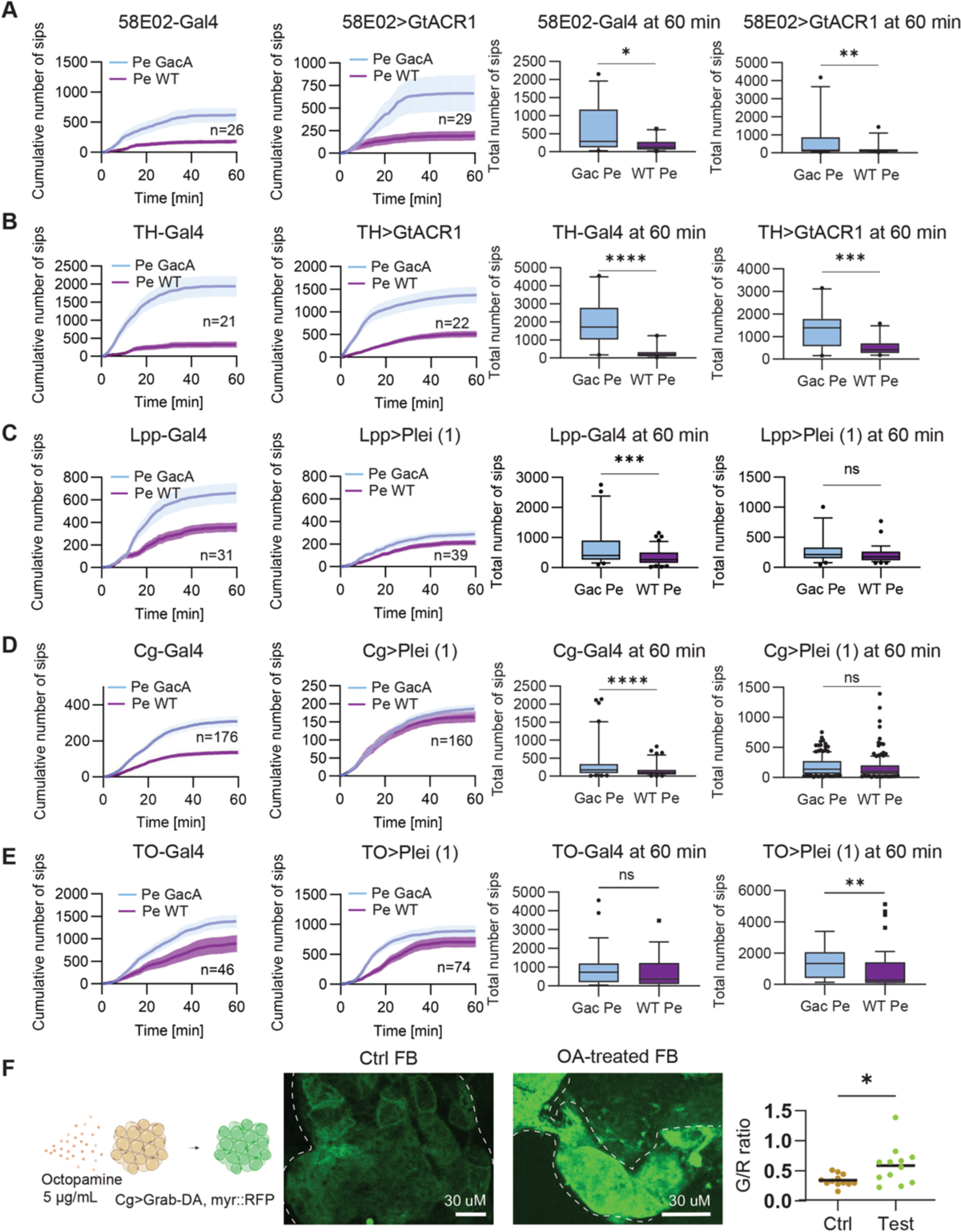
Tyrosine hydroxylase is required in the fat body for reducing pathogen consumption. (A) Feeding preferences for harmless and harmful *Pe* bacteria upon optogenetic inhibition of PAM cluster DANs using 58E02-Gal4;UAS-GtACR1 (n = 29), and control flies 58E02-Gal4 (n = 26). Cumulative number of sips; mean ± SEM. Total number of sips at 60 mins; *p-values* calculated via Wilcoxon matched-pairs signed rank test. (B) Feeding preferences for harmless and harmful *Pe* bacteria upon optogenetic inhibition of PPL1 cluster DANs using TH-Gal4;UAS-GtACR1 (n = 22), and control flies TH-Gal4 (n = 21). Cumulative number of sips; mean ± SEM. Total number of sips at 60 mins; *p-values* calculated via Wilcoxon matched-pairs signed rank test. (C) Feeding preferences for harmless and harmful *Pe* bacteria upon knock-down of *pale* (tyrosine hydroxylase) in fat body using LPP-Gal4;UAS-Ple RNAi (n=39), and control flies LPP-Gal4 (n = 31). Cumulative number of sips; mean ± SEM. Total number of sips at 60 mins; *p-values* calculated via Wilcoxon matched-pairs signed rank test. (D) Feeding preferences for harmless and harmful *Pe* bacteria upon knock-down of tyrosine hydroxylase in fat body using the alternative fat body driver Cg-Gal4;UAS-Ple RNAi (n = 160), and control flies Cg-Gal4 (n = 176). Cumulative number of sips; mean ± SEM. Total number of sips at 60 mins; *p-values* calculated via Wilcoxon matched-pairs signed rank test. (E) Feeding preferences for harmless and harmful *Pe* bacteria upon knock-down of tyrosine hydroxylase in the head fat body using TO-Gal4;UAS-Ple RNAi (n = 74), and control flies TO-Gal4 (n = 46). Cumulative number of sips; mean ± SEM. Total number of sips at 60 mins; *p-values* calculated via Wilcoxon matched-pairs signed rank test. (F) Representative maximum intensity Z-projection images of the fat body expressing Grab-DA with or without OA stimulation. Green fluorescence quantified as G (green)/R (red) ratio was significantly higher in the OA-exposed group (n = 12) compared to the control (saline) group (n = 12). *p-values* were calculated using the Welch-corrected t-test. For all analyses, the statistical notation is as follows: ns (not significant), p > 0.05; * p < 0.05; ** p < 0.01; *** p < 0.001; **** p < 0.0001. Error bars in all panels represent the standard error of the mean (SEM).

However, since the gene *ple* was enriched in the fat body tissue upon harmful *Pe* ingestion, we hypothesized that dopamine could be produced by the fat body in response to pathogen infection and contribute to the observed behavioral adaptation. We further hypothesized that the octopaminergic neuron-induced increase in calcium in fat cells could trigger dopamine release from the fat body reminiscent to the situation in dopaminergic neurons upon activation. To test this, we downregulated *ple* in the fat body again by RNA interference. Remarkably, we observed that flies with reduced levels of TH in their fat body fed equally on both pathogenic and harmless *Pe* (Fig. 4C, S4D). We validated this finding by using an alternative fat body driver and two independent *ple* RNAi lines (Fig. 4D, S4C). In addition, we also knocked down *ple* exclusively in the head fat body and observed that the flies did not avoid feeding on pathogen-laced food (Fig. 4E). As one of the drivers, Cg-Gal4 also drives expression in the hemocytes, and a recent paper suggested that detection of Gram-negative bacteria promotes hematopoiesis ^51^, we also downregulated *ple* in hemocytes selectively through a hemocyte specific Gal4-driver. However, flies with less TH in hemocytes retained their ability to avoid feeding from harmful bacteria-contaminated food (Fig. S4G).

To gain more direct evidence for a release of dopamine by the fat body, we expressed the genetically encoded dopamine sensor GRAB-DA in the fat body (Fig. 4F) ^52^. In addition to GRAB-DA, we co-expressed mRFP to normalize for the size of the fat body to allow for quantification of a putative increase in GRAB-DA fluorescence. We again incubated explant fat body tissue with octopamine immediately prior to imaging. Our results indicated that stimulation with octopamine not only increased intracellular calcium levels (see Fig. 3D,E) but also stimulated dopamine release from the fat body (Fig. 4F).

Taken together, our results provide evidence that dopamine release from the fat body in response to octopaminergic stimulation enables flies to adapt their behavior to an oral infection by suppressing their appetite for harmful bacteria.

### Dopamine receptor-mediated feeding choice acts in Mushroom body output neurons

Dopamine functions primarily via activating dopamine receptors ^53^. Flies possess two distinct types of dopamine receptors: the D1-like receptors, specifically Dop1R1 and Dop1R2, and the D2-like receptor known as Dop2R ^54^. Consequently, we proceeded to investigate the potential involvement of dopamine receptors in the observed avoidance of hazardous *Pe*. Indeed, flies that lack Dop1R1 no longer expressed a preference for harmless over pathogenic *Pe* (Fig. 5A) indicating that Dop1R1 is required for post-ingestion pathogen feeding avoidance. By contrast, the *Dop2R* mutant flies exhibited no discernible difference when compared to control flies (Fig. 5B).

**Figure 5.**
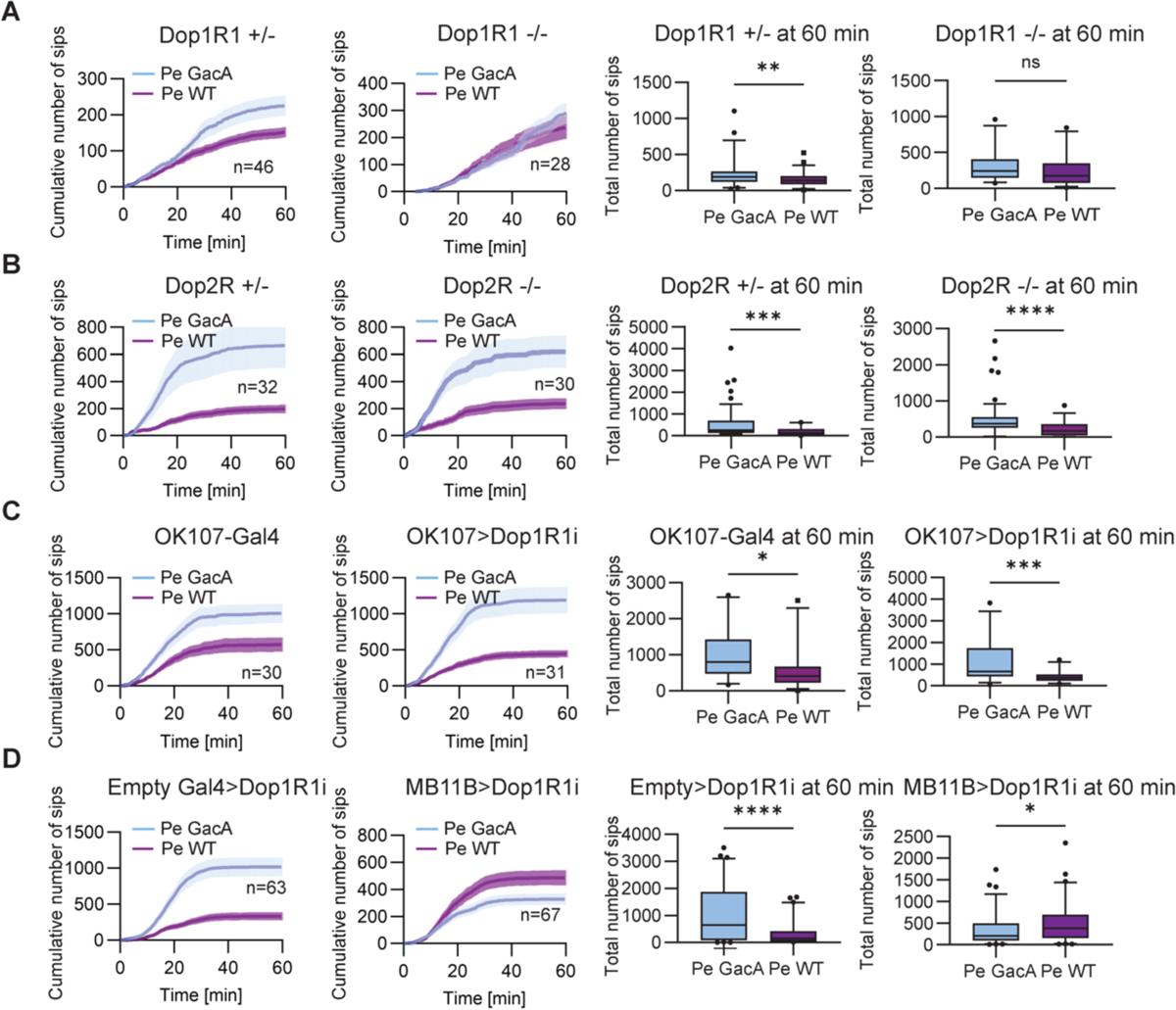
Dopamine receptor Dop1R1 in specific MBONs regulates pathogen aversion. (A) Feeding preferences for harmless and harmful *Pe* bacteria for homozygous mutant Dop1R1 flies (Dop1R1-/-) and heterozygous control flies: n = 28 (Dop1R1-/-), n = 46 (Dop1R1+/-). Cumulative number of sips; mean ± SEM. Total number of sips at 60 mins; *p-values* calculated via Wilcoxon matched-pairs signed rank test. (B) Feeding preferences for harmless and harmful *Pe* bacteria for homozygous mutant Dop2R flies (Dop2R-/-) and heterozygous control flies: n = 30 (Dop2R-/-), n = 32 (Dop2R+/-). Cumulative number of sips; mean ± SEM. Total number of sips at 60 mins; *p-values* calculated via Wilcoxon matched-pairs signed rank test. (C) Feeding preferences for harmless and harmful *Pe* bacteria upon knock-down of Dop1R1 in all MB Kenyon cells using OK107-Gal4;UAS-Dop1R1 RNAi (n = 31) and control flies OK107-Gal4 (n = 30). Cumulative number of sips; mean ± SEM. Total number of sips at 60 mins; *p-values* calculated via Wilcoxon matched-pairs signed rank test. (D) Feeding preferences for harmless and harmful *Pe* bacteria upon knock-down of Dop1R1 in MBON 01, 03 and 04 (aka γ5β’2α, β’2mp, and β’2mp_bilateral MBON) using MB11B-Gal4;UAS-Dop1R1 RNAi (n = 67) and control flies (empty driver pBDPU-Gal4;UAS-Dop1R1 RNAi (n = 63). Cumulative number of sips; mean ± SEM. Total number of sips at 60 mins; *p-values* calculated via Wilcoxon matched-pairs signed rank test. For all analyses, the statistical notation is as follows: ns (not significant), p > 0.05; * p < 0.05; ** p < 0.01; *** p < 0.001; **** p < 0.0001. Error bars in all panels represent the standard error of the mean (SEM).

Therefore, we asked where Dop1R1 is required for pathogen feeding avoidance. Classical associative olfactory learning demonstrated that Dop1R1 is essential in the MB Kenyon cells (KCs) for aversive and appetitive learning in *Drosophila* ^55,56^. However, we downregulated Dop1R1 expression in KCs and found that flies with reduced Dop1R1 expression in the KCs could still differentiate between harmless and pathogenic forms of the bacteria (Fig. 5C).

Our prior work has shown that a subset of MBONs (MBON01, 03, 04) is required for the post-ingestion avoidance of harmful bacteria ^30^. Given the involvement of these specific MBONs, we knocked down Dop1R1 expression in these MBONs using a specific split-Gal4 driver ^57^. Remarkably, these flies did not only lose the ability to suppress feeding on harmful bacteria, but even preferred feeding on pathogenic *Pe* bacteria over the harmless strain, consistent with the flies’ innate, pre-ingestion, preference for harmful *Pe* and to the phenotype observed in flies lacking most AMPs (Fig. 5D; see Fig. 1B).

Taken together, our findings show that Dop1R1 in MBONs is required for post-ingestion avoidance of pathogen-containing food. Based on these data and our findings above, we propose that dopamine, released by the fat body upon bacterial infection, acts as a ligand for Dop1R1 in MBONs and, thereby, reduces the fly’s preference for a dangerous pathogen.

## Discussion

In this study, we provide evidence of a direct crosstalk between the immune and metabolic organ fat body and neuromodulatory OANs in response to pathogen ingestion in *Drosophila melanogaster*. Upon detection of pathogenic bacteria via the innate immune receptors PGRPs, both the fat body and OANs release AMPs, such as DptB, presumably to fight and limit the spread of the infection. Importantly, OANs innervate the fat body where they release octopamine, which activates octopamine receptor signaling and in turn triggers an increase of calcium and the release of dopamine from fat cells. Dopamine through Dop1R1 in specific MBONs modulates food preference leading to the lasting reduction of pathogen ingestion (Fig. 6). These findings reveal a complex interaction between the immune system, metabolism and the nervous system in insects, providing new insights into how other organs besides the brain can contribute to modulating behavior and enhance the organism’s survival chances when faced with infection.

**Figure 6.**
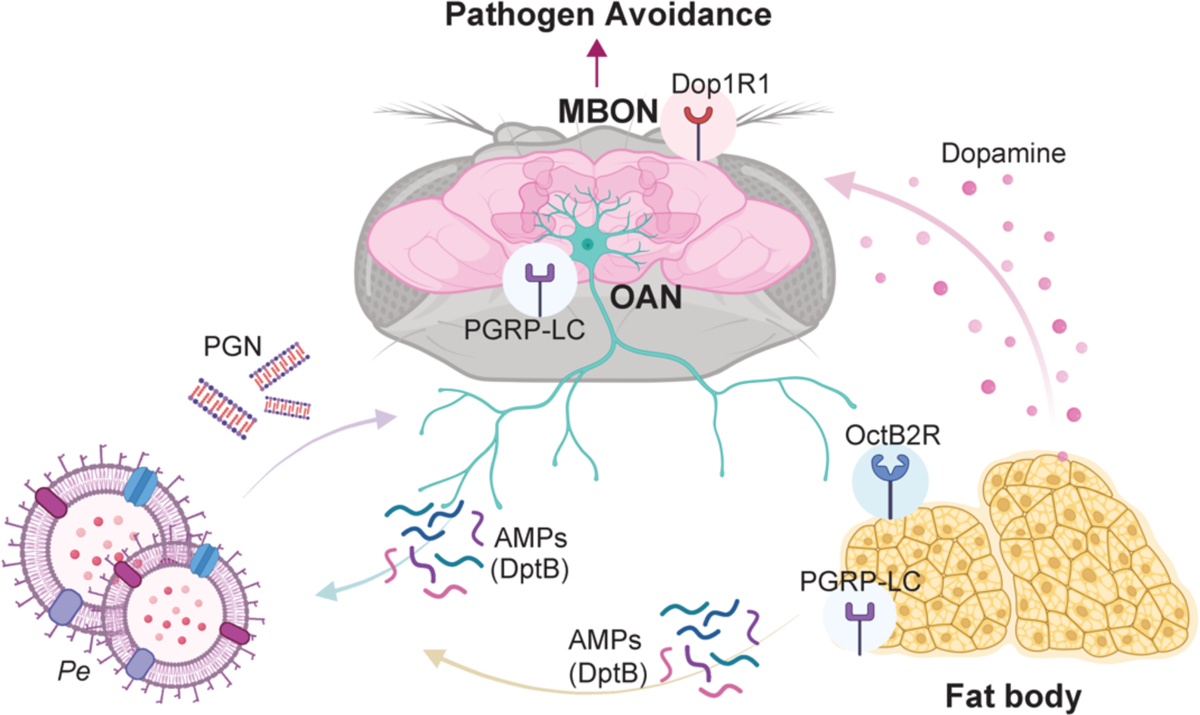
A model for a nervous system fat body axis in post-ingestion pathogen aversion. Pathogen ingestion through food is detected via PGRP-LC innate immune receptors in octopaminergic neurons (OAN) and the fat bodies. This leads to the production of antimicrobial peptides (AMPs), i.e. DiptericinB (DptB), to combat the infection. PGRP-LC activation also leads to the activation of OANs and to the subsequent stimulation of fat body cells via Octβ2R signaling in fat cells. This induces production and release of dopamine from the fat body. The released dopamine then signals to the mushroom body (MB) and specific MB output neurons (MBONs) leading to a significant reduction in pathogen ingestion.

### Why does post-ingestion pathogen avoidance develop within minutes?

The anatomical proximity of the head fat body and OANs, of which many innervate neuropils close to the esophagus, may contribute to their function in adaptive behavior (see Fig. 3). It is conceivable that PGRPs from bacteria are, therefore, first sensed by the head fat body and OAN clusters around the esophagus. Such an architecture could significantly decrease the latency of pathogen detection and explain the rapid onset of subsequent behavioral changes. Despite multiple attempts, we have not been able to pinpoint the specific cluster of octopaminergic neurons responsible for this behavior due to the lack of suitable genetic tools and Gal4-driver lines. Identifying these neurons will remain a focus of our future studies. We have, however, identified an octopamine receptor in the fat body critical for post-ingestion pathogen avoidance. Octβ2R is expressed in the fat body and its removal specifically from the head fat body was sufficient to prevent the flies from choosing food laced with harmless bacteria over pathogen-contaminated food (see Fig. 3). It is possible that OAMB or Octα2R are involved as well. Lethality and the lack of an RNA interference phenotype, respectively, prevented us from experimentally testing this possibility. Nevertheless, the involvement of the head fat body and its position close to the esophagus and OANs might explain the rapid adaptation of food choice behavior. Interestingly, such a rapid detection during food consumption might help to prime the immune system and improve the efficacy of the immune response as suggested by recent data in the fly larva ^51^. It will be interesting to investigate how an octopamine triggered activation of fat cells through calcium signals boosts dopamine release as well as the production and release of AMPs.

### How does fat body dopamine interact with neurons in the brain to change behavior?

Our data provide significant evidence for the production and release of dopamine from the fat body upon pathogen infection. Notably, dopamine from DANs does not play a role in this process as optogenetic inhibition or knock-down of tyrosine hydroxylase (aka ple) did not induce a phenotype, consistent with our earlier findings (see Fig. 4) ^30^. In addition, we show that stimulation with octopamine or optogenetic activation of OANs induces an increase in calcium levels in the fat body (see Fig. 3 and 4). While we do not know yet, how dopamine release is regulated in the fat body and how dopamine travels to the brain, passes the hemolymph-brain-barrier and interacts with Dop1R1 in MBONs, previous work supports the hypothesis that this is possible ^15^. The authors showed that fat tissue in humans, mice and flies expresses components of the dopamine release machinery (e.g. the vesicular monoamine transporter Vmat). Moreover, the dopamine release cycle is increased in fat cells of severely obese humans indicating that fat tissue can adjust its release of dopamine in pathological conditions ^15^. Association studies in humans and genetic experiments in *Drosophila* further prompted the authors to propose a direct impact of fat tissue on cognitive abilities. More specifically, RNAi-mediated inhibition of fat body expressed neuronal genes as well as overexpression of release cycle–associated proteins SLC18A2 and RIMS1 promoted courtship learning in flies and modulated the expression of learning-associated genes such as the dunce (aka cAMP-phosphodiesterase) and rutabaga (aka calcium-sensitive dependent adenylate cyclase). These data support our findings and conclusion of a connection between fat body and brain function ^15^. While dopamine in mammals generally does not cross the blood-brain-barrier (BBB) efficiently ^58^, it is not known whether the same is true for the fly’s BBB. Moreover, the mammalian BBB is leaky in the region of the arcuate nucleus and median eminence to allow communication with the body and entry of peripheral signals such as leptin, ghrelin and nutrients ^59^. Notably, Vmat is expressed in the glia cells that form the fly’s BBB and might promote the transport of dopamine ^15^. How dopamine crosses the BBB is an important question for future work.

To reduce pathogen-ingestion, dopamine appears to act through Dop1R1 in specific MBONs (MBON01, 03, and 04; see Fig. 5). Previous work including ours has shown that MBONs providing output from the β2, β’2 and γ5-compartments of the MB promote innate and learned avoidance when activated by odor or optogenetics ^57,60–62^. We have further shown in Kobler et al., 2020, that post-ingestion avoidance of pathogenic food required a functional olfactory system as flies lacking the olfactory receptor ORCO no longer avoid pathogenic bacteria upon ingestion ^30^. Given that Dop1R1 signaling through activation of adenylyl cyclase promotes neuronal activation, we propose that the MB and these MBONs act as integrators of pathogen odor and post-ingestion immune responses. In this scenario, dopamine, released upon ingestion of pathogenic bacteria, enhances the activation of these MBONs by bacterial odor and thereby promotes MBON-dependent avoidance and reduces further pathogen ingestion.

### Which role do AMPs play in regulating pathogen avoidance behavior?

Our data and previous publications suggest that PGRP signaling plays versatile roles in the nervous system (see introduction). However, we still lack a full understanding of how it contributes to the adaptation of behavior. We have shown that OAN-triggered activation of the fat body and subsequent release and action of dopamine suppresses pathogen feeding quickly after initial ingestion. An additional mechanism for this adaptation of behavior could be that AMPs may function as signaling peptides akin to neuropeptides. Flies lacking most AMPs (ΔAMP mutants) prefer pathogenic bacteria over harmless controls even after 1h of feeding (see Fig. 1). Interestingly, AMPs exhibit numerous similarities to neuropeptides, including size, cationic charge, amphipathic structure, and amino acid content. Moreover, AMPs can be found in the central nervous system of various species ^6,63–65^. In *C. elegans*, a group of genes called *nlp* produces multiple infection-induced proteins known as NLPs. These proteins are expressed in neurons and are believed to have both neuromodulatory and antimicrobial functions. For instance, NLP-29 regulates dendrite degeneration through neuropeptide receptor 12 ^66–68^. In humans, pituitary adenylate cyclase-activating polypeptide (PACAP) exhibits antibacterial properties and is found in the central nervous system, where it acts as a neuropeptide ^69^. Notably, within minutes of an infection, AMPs can be detected in the fly’s head and hemolymph ^70^. This timeframe is comparable to the time required for flies to adapt their behavior and cease feeding on pathogen-contaminated food (see Fig. 1). Finally, the change in feeding behavior we observed in our feeding assays in flies with reduced levels of AMPs in the fat body is also consistent with a more general role of AMPs in appetite regulation (see Fig. 1). In humans, the liver-expressed antimicrobial peptide-2 (LEAP2) enhances appetite and food intake by inhibiting the growth hormone secretagogue receptor (GHSR), aka ghrelin receptor ^71^. Although we have not identified a neural receptor for AMPs in insects up to now, it is plausible that AMPs are involved as direct signals of infection underpinning nervous-system mediated changes of feeding behavior and appetite.

### Concluding remarks

Our data reveal a bidirectional signaling mechanism between nervous system and fat body through neuronal innervation of adipose tissue, on the one hand, and the systemic action of fat body-secreted neuromodulators on neurons, on the other hand. Given the danger pathogen ingestion poses on the entire organism, we propose that the systemic action of a neuromodulator such as dopamine, as opposed to local release by specific neurons, can trigger the adaptation of multiple survival-related behaviors, including feeding and reproduction, efficiently and in sync. This situation might be comparable to the combined systemic and local action of adrenaline and noradrenaline, respectively, in the sympathetic nervous system in stressful or dangerous situations. Of note, octopamine is thought to be the insect counterpart of noradrenaline and dopamine is the essential precursor in the production of noradrenaline. Thus, given the role of sympathetic innervation of fat and the liver through noradrenergic neurons in the regulation of metabolism, thermogenesis, immunity and adipocyte differentiation in mammals ^72,73^, the here described mechanism in insects might have evolved from a common origin. The essential role of innate immune receptors in the presented scenario in insects might explain the close, at times pathological, relationship between immune system, nervous system and metabolic organs observed in humans. Findings in the fly might, thus, help to accelerate our conceptual and mechanistic understanding of the important neuro-immuno-metabolism axis in health and disease.

## Author contributions

I.C.G.K. and Y.W. conceptualized the study and designed the experiments. Y.W. carried out and analyzed all behavioral experiments, genetic manipulations, histological and imaging experiments. K.S. and H.T. carried out the expression analysis of different octopamine receptors in different tissues. I.C.G.K. and Y.W. wrote the manuscript with the help of all authors.

## Acknowledgments

We thank Heidi Miller-Mommerskamp and Natalie Lindenberg for help in general fly husbandry and bacterial work. We also wish to thank Jean-François De Backer for his help with all histological and live imaging experiments. We acknowledge Johanna Kobler for training in all behavioral analysis and bacterial work, and we thank Bruno Lemaitre and Mark Hanson for providing the reagents to test the contribution of AMPs and their generous advice on immunity-related experiments and methods. We are grateful to Nicolas Gompel, Ana Domingos, Aurélie Muria, and Jean-François De Backer for their thoughtful comments on the manuscript. This project was generously funded by the German Research Foundation (FOR2705/TP3, GR4310/6-2 to IGK; INST 217/1135-1 to IGK).

## Supplemental information

Figures S1-S4

Table S1

## Methods

### EXPERIMENTAL MODEL AND SUBJECT DETAILS

#### Flies strains and maintenance

Flies were raised under a 12:12 hours light/dark cycle on standard cornmeal food at 25°C and 60% humidity. For starvation experiments, flies were transferred to a starvation vial containing a wet tissue paper as water source 24 hours prior experiments. For optogenetic experiments, adult flies were collected after hatching and kept on all trans-retinal supplemented food (1:250) under blue light only conditions. All experiments were conducted on 5-8-day old female flies, unless stated elsewhere.

## METHOD DETAILS

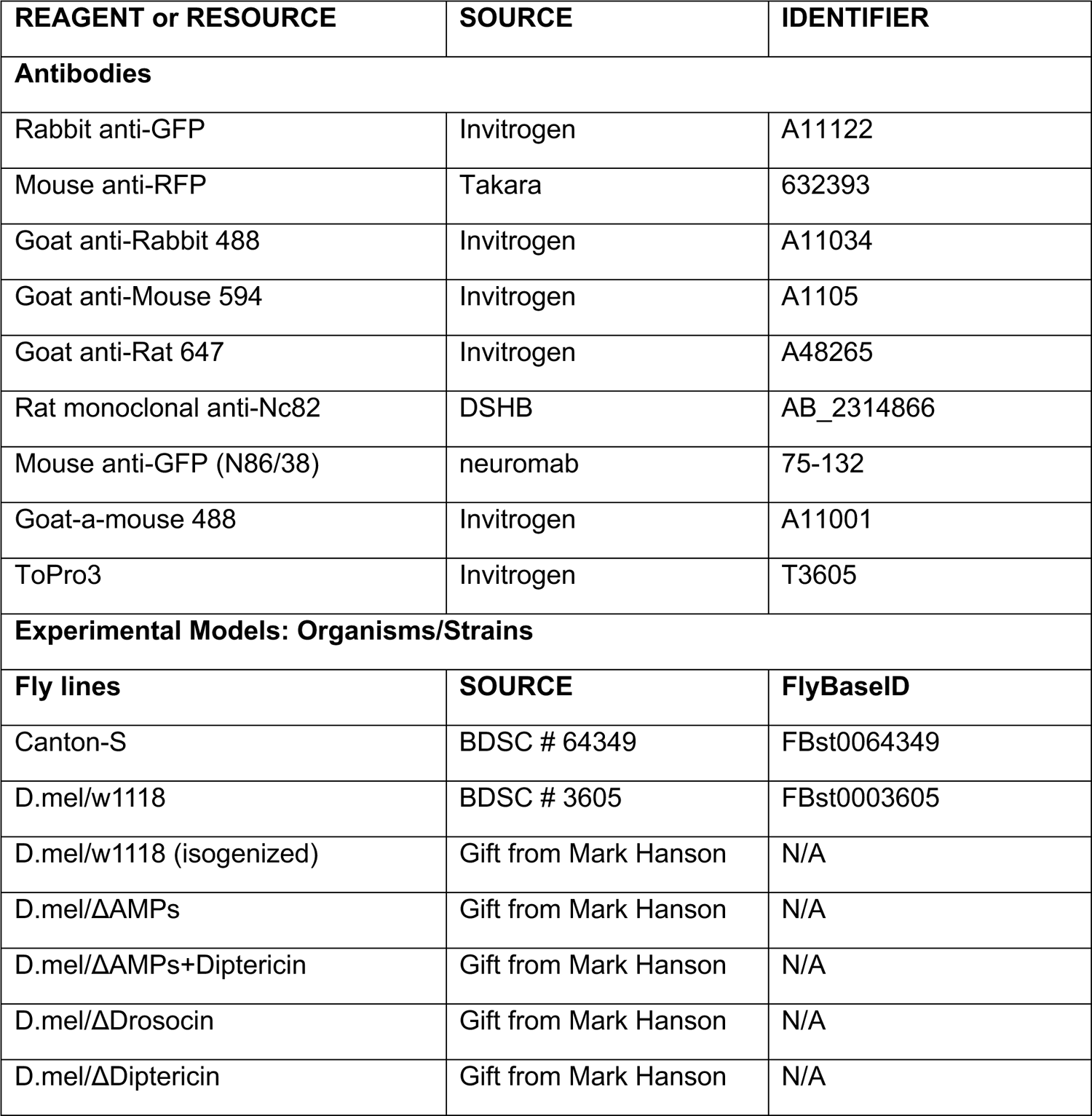
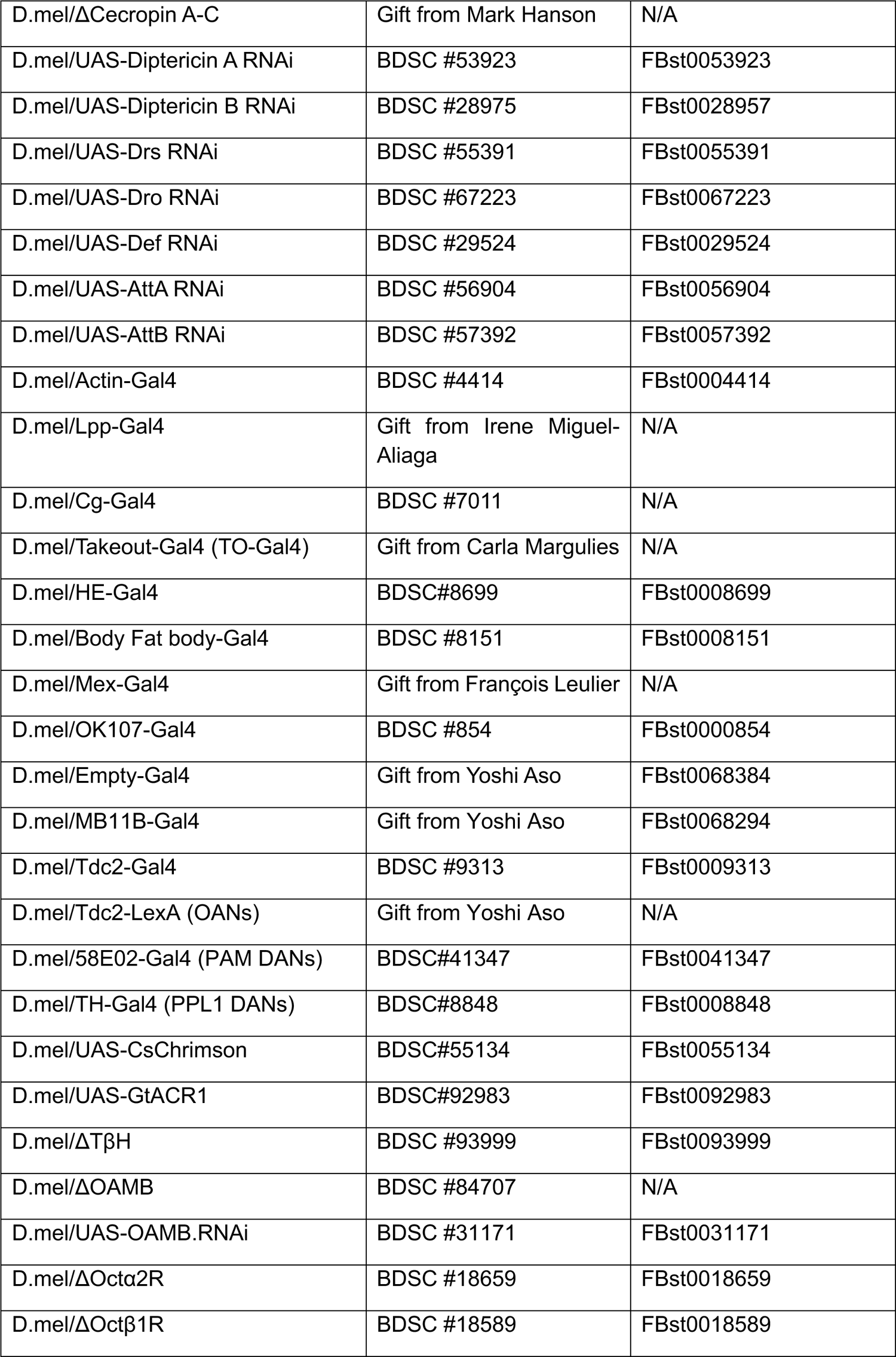
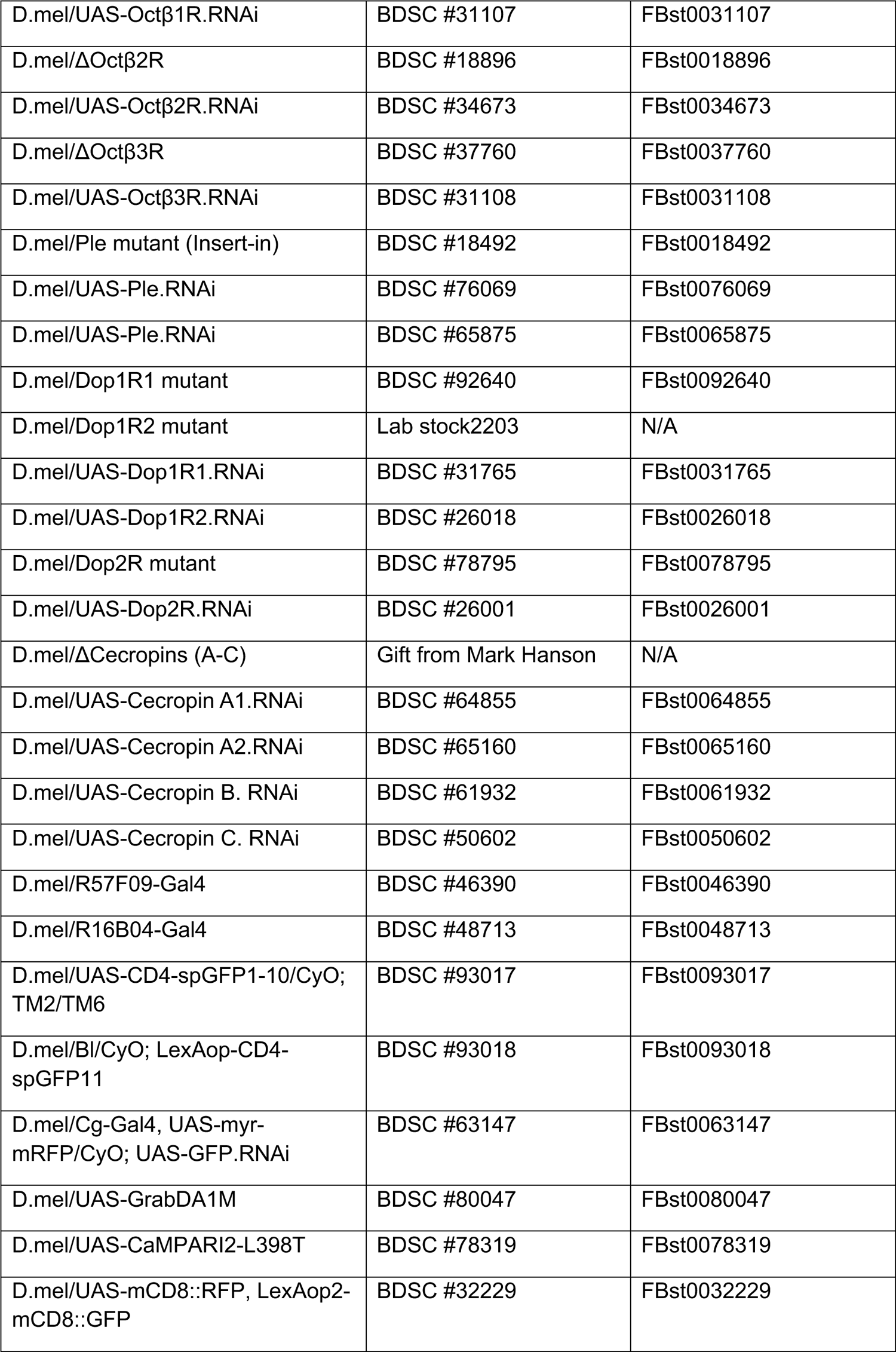
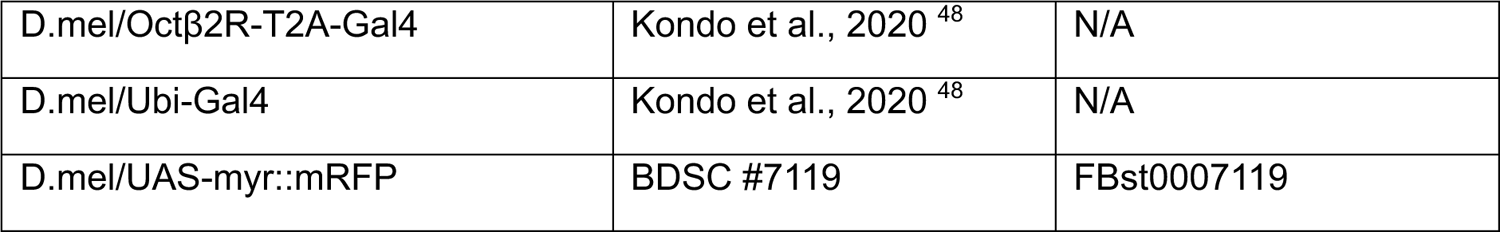

| REAGENT or RESOURCE | SOURCE | IDENTIFIER |
| --- | --- | --- |
| **Antibodies** |  |  |
| Rabbit anti-GFP | Invitrogen | A11122 |
| Mouse anti-RFP | Takara | 632393 |
| Goat anti-Rabbit 488 | Invitrogen | A11034 |
| Goat anti-Mouse 594 | Invitrogen | A1105 |
| Goat anti-Rat 647 | Invitrogen | A48265 |
| Rat monoclonal anti-Nc82 | DSHB | AB_2314866 |
| Mouse anti-GFP (N86/38) | neuromab | 75-132 |
| Goat-a-mouse 488 | Invitrogen | A11001 |
| ToPro3 | Invitrogen | T3605 |
| **Experimental Models: Organisms/Strains** |  |  |
| **Fly lines** | **SOURCE** | **FlyBaseID** |
| Canton-S | BDSC # 64349 | FBst0064349 |
| D.mel/w1118 | BDSC # 3605 | FBst0003605 |
| D.mel/w1118 (isogenized) | Gift from Mark Hanson | N/A |
| D.mel/ΔAMPs | Gift from Mark Hanson | N/A |
| D.mel/ΔAMPs+Diptericin | Gift from Mark Hanson | N/A |
| D.mel/ΔDrosocin | Gift from Mark Hanson | N/A |
| D.mel/ΔDiptericin | Gift from Mark Hanson | N/A |
| D.mel/ΔCecropin A-C | Gift from Mark Hanson | N/A |
| D.mel/UAS-Diptericin A RNAi | BDSC #53923 | FBst0053923 |
| D.mel/UAS-Diptericin B RNAi | BDSC #28975 | FBst0028957 |
| D.mel/UAS-Drs RNAi | BDSC #55391 | FBst0055391 |
| D.mel/UAS-Dro RNAi | BDSC #67223 | FBst0067223 |
| D.mel/UAS-Def RNAi | BDSC #29524 | FBst0029524 |
| D.mel/UAS-AttA RNAi | BDSC #56904 | FBst0056904 |
| D.mel/UAS-AttB RNAi | BDSC #57392 | FBst0057392 |
| D.mel/Actin-Gal4 | BDSC #4414 | FBst0004414 |
| D.mel/Lpp-Gal4 | Gift from Irene Miguel-Aliaga | N/A |
| D.mel/Cg-Gal4 | BDSC #7011 | N/A |
| D.mel/Takeout-Gal4 (TO-Gal4) | Gift from Carla Margulies | N/A |
| D.mel/HE-Gal4 | BDSC#8699 | FBst0008699 |
| D.mel/Body Fat body-Gal4 | BDSC #8151 | FBst0008151 |
| D.mel/Mex-Gal4 | Gift from François Leulier | N/A |
| D.mel/OK107-Gal4 | BDSC #854 | FBst0000854 |
| D.mel/Empty-Gal4 | Gift from Yoshi Aso | FBst0068384 |
| D.mel/MB11B-Gal4 | Gift from Yoshi Aso | FBst0068294 |
| D.mel/Tdc2-Gal4 | BDSC #9313 | FBst0009313 |
| D.mel/Tdc2-LexA (OANs) | Gift from Yoshi Aso | N/A |
| D.mel/58E02-Gal4 (PAM DANs) | BDSC#41347 | FBst0041347 |
| D.mel/TH-Gal4 (PPL1 DANs) | BDSC#8848 | FBst0008848 |
| D.mel/UAS-CsChrimson | BDSC#55134 | FBst0055134 |
| D.mel/UAS-GtACR1 | BDSC#92983 | FBst0092983 |
| D.mel/ΔTβH | BDSC #93999 | FBst0093999 |
| D.mel/ΔOAMB | BDSC #84707 | N/A |
| D.mel/UAS-OAMB.RNAi | BDSC #31171 | FBst0031171 |
| D.mel/ΔOctα2R | BDSC #18659 | FBst0018659 |
| D.mel/ΔOctβ1R | BDSC #18589 | FBst0018589 |
| D.mel/UAS-Octβ1R.RNAi | BDSC #31107 | FBst0031107 |
| D.mel/ΔOctβ2R | BDSC #18896 | FBst0018896 |
| D.mel/UAS-Octβ2R.RNAi | BDSC #34673 | FBst0034673 |
| D.mel/ΔOctβ3R | BDSC #37760 | FBst0037760 |
| D.mel/UAS-Octβ3R.RNAi | BDSC #31108 | FBst0031108 |
| D.mel/Ple mutant (Insert-in) | BDSC #18492 | FBst0018492 |
| D.mel/UAS-Ple.RNAi | BDSC #76069 | FBst0076069 |
| D.mel/UAS-Ple.RNAi | BDSC #65875 | FBst0065875 |
| D.mel/Dop1R1 mutant | BDSC #92640 | FBst0092640 |
| D.mel/Dop1R2 mutant | Lab stock2203 | N/A |
| D.mel/UAS-Dop1R1.RNAi | BDSC #31765 | FBst0031765 |
| D.mel/UAS-Dop1R2.RNAi | BDSC #26018 | FBst0026018 |
| D.mel/Dop2R mutant | BDSC #78795 | FBst0078795 |
| D.mel/UAS-Dop2R.RNAi | BDSC #26001 | FBst0026001 |
| D.mel/ΔCecropins (A-C) | Gift from Mark Hanson | N/A |
| D.mel/UAS-Cecropin A1.RNAi | BDSC #64855 | FBst0064855 |
| D.mel/UAS-Cecropin A2.RNAi | BDSC #65160 | FBst0065160 |
| D.mel/UAS-Cecropin B. RNAi | BDSC #61932 | FBst0061932 |
| D.mel/UAS-Cecropin C. RNAi | BDSC #50602 | FBst0050602 |
| D.mel/R57F09-Gal4 | BDSC #46390 | FBst0046390 |
| D.mel/R16B04-Gal4 | BDSC #48713 | FBst0048713 |
| D.mel/UAS-CD4-spGFP1-10/CyO;<br>TM2/TM6 | BDSC #93017 | FBst0093017 |
| D.mel/BI/CyO; LexAop-CD4-spGFP11 | BDSC #93018 | FBst0093018 |
| D.mel/Cg-Gal4, UAS-myr-mRFP/CyO; UAS-GFP.RNAi | BDSC #63147 | FBst0063147 |
| D.mel/UAS-GrabDA1M | BDSC #80047 | FBst0080047 |
| D.mel/UAS-CaMPARI2-L398T | BDSC #78319 | FBst0078319 |
| D.mel/UAS-mCD8::RFP, LexAop2-mCD8::GFP | BDSC #32229 | FBst0032229 |
| D.mel/Octβ2R-T2A-Gal4 | Kondo et al., 2020<sup>48</sup> | N/A |
| D.mel/Ubi-Gal4 | Kondo et al., 2020<sup>48</sup> | N/A |
| D.mel/UAS-myr::mRFP | BDSC #7119 | FBst0007119 |

### Bacterial strains and preparation

The Gram-negative bacterium *Pseudomonas entomophila* (*Pe*), including both the wild-type (pathogenic) and *gacA* (avirulent) strains, and also *Erwinia carotovora carotovora 15* (*Ecc15*) strains *Ecc15 pOM1* (pathogenic) and *Ecc15 evf* (avirulent, rifampicin-resistant) were generously provided by Bruno Lemaitre and used for infecting flies. The *Pe* strains were inoculated onto lysogeny broth (LB) agar plates plates (composed of 10 g NaCl, 10 g tryptone, 5 g yeast, and 15 g agar per 1 L) added with 100 μg/ml rifampicin and 1% skim milk (Sigma-Aldrich #70166). The plates were incubated at a temperature of 30 for a minimum of 24 hours for *Pe WT* or 30 hours for *Pe gacA*. Skim milk was employed as a marker to ascertain pathogenicity. Pathogenic clones of *Pe WT* exhibited proteolytic activity, leading to the development of transparent colonies on the otherwise hazy LB/milk agar plates. However, *Pe gacA* lacked this protease, resulting in the creation of plates with a blurry appearance. The *Ecc15* strains were grown on LB agar plates with 100 μg/ml antibiotics respectively (spectinomycin for *Ecc15 pOM1,* rifampicin for *Ecc15 evf)* at 29°C for 24 hours. The bacteria were inoculated weekly and maintained at a temperature of 4°C.

In preparation for experiments, liquid cultures have been used to produce a substantial amount of bacteria: either a protease-positive *Pe WT* clone (clear colony) or a single colony of *Pe gacA* was pre-cultured in 20 ml of LB medium with 100 μg/ml rifampicin. The pre-cultures were incubated for 8 hours at 250 rpm and a temperature of 30°C. Afterward, an overnight culture was developed by diluting the pre-culture in LB medium with 100 μg/ml rifampicin at a ratio of 1:16. The culture was thereafter cultivated for at least 16 hours at a speed of 250 rpm at a temperature of 30°C. For *Ecc15* strains, single colonies of *Ecc15 evf* and *Ecc15 pOM1* were directly cultured in LB medium containing 100 μg/ml respective antibiotic overnight at 29°C with shaking speed at 220 rpm.

To obtain a concentrated bacterial pellet, the bacterial cultures were centrifuged at a force of 2500 g for 15 minutes at a temperature of 4°C, after measuring the optical density at a wavelength of 600 nanometers (nm) (OD600). Following that, the liquid component above the solid sediment was mostly removed, while the solid pellets were immediately blended with the remaining medium. The concentrated bacterial culture was adjusted to achieve the desired optical density (OD600) of approximately 200 using PBS. Subsequently, it was maintained at a temperature of 4°C for a period of 2 hours. before being utilized.

### Bacterial infection through ingestion

The present study examined the behavioral reactions triggered by the ingestion of both pathogenic and non-pathogenic bacteria. Consequently, flies had to consume microbes through feeding. To motivate flies to feed, they were subjected to a period of dry starvation in empty bottles for 3 to 5 hours at a temperature of 25°C.

### Behavioral assays

#### FlyPAD and OptoPAD food choice experiments

FlyPAD food choice experiments were conducted as described previously ^30^. Individual flies were transferred by aspiration into a behavioral arena, where they were allowed to feed on two gelled food substrates for one hour. The tested food choices were Pe GacA + 5% sucrose versus Pe WT + 5% sucrose, with all substrates containing 1% agarose (low gelling temperature, A9414, Sigma-Aldrich). The experiments were carried out in a climate-controlled chamber at 25°C and 60% humidity. For optogenetic experiments, we used an open-loop stimulation paradigm. Here, GtACR1 was activated by continuous green light (523 nm, 140 µW/mm²) for 1 hour, starting when the fly’s proboscis first contacted the sucrose. Capacitance signal analysis was performed using a custom MATLAB script provided by Pavel Itskov ^37,74^, and feeding behavior was evaluated by summing the cumulative and total number of sips from the two electrodes.

#### Tissue-specific RNAseq at different time course upon pathogen ingestion

The present study examined the changes in gene expression across various tissues and temporal periods after the oral consumption of pathogenic bacteria Pe WT. Thus, this experiment required the flies to consume Pe WT through oral ingestion. To boost the flies’ motivation to consume the harmful bacteria, they were deprived of food before the experiments, and the bacteria was combined with sucrose.

As stated in the section on bacteria preparation for feeding assay, the bacterial solution with an optical density (OD) of approximately 600 was mixed in equal proportions with a 10% sucrose solution. As a result, the final concentration of OD600 was approximately 100 and the sucrose content was 5%. Afterwards, the flies were placed into a standard fly bottle with 1.5% agarose, which served to maintain an ideal level of humidity. Furthermore, a filter paper soaked in a solution containing bacteria and sucrose mixture. The control treatment in our experiment was using a 5% sucrose solution in PBS as the PBS was used to dilute the bacteria pellet.

#### Tissue preparation and RNA extraction

The experiment employed wild-type Canton-S flies in all cases. Female flies aged 4-7 days were collected and subjected to starvation for 3-5 hours. Subsequently, they were divided into standard fly bottles and fed with bacteria or controls.

After that, each fly was euthanized in ethanol and promptly dissected in ice-cold RNAlater. The heads, guts, and abdomen fat bodies were swiftly collected and individually placed in three 200 μl microcentrifuge tubes containing RNAlater on ice. The tubes were then stored at −20C immediately till RNA extraction. Each condition had three biological replicates. The tissues in each tube were homogenized using a Beads homogenizer. The process of extracting and purifying RNA was carried out using the RNeasy mini kit from QIAGEN. The RNA was extracted using sterile water, and its purity and concentration were assessed by measuring the 260/280 and 260/230 ratios using a NanoDrop 2000c (Thermo Scientific).

#### mRNA-seq libraries and sequencing

The purity of the total RNA from twenty-seven RNA samples, comprising of three duplicates of three tissues from each of the three feeding groups, was verified using the Qubit RNA Broad Range Assay (Invitrogen). 200 ng of total RNA was used for library preparation for each sample. The initial stage of the library preparation process entailed the amplification of cDNA using poly-dT primers, obviating the requirement for mRNA purification utilizing beads.

The libraries were prepared using the 3’mRNA-Seq Library Prep Kit FWD with Unique Dual Indices (Lexogen; User guide 113UG227V0100) following the manufacturer’s instructions. The Qubit dsDNA High Sensitivity Assay (Invitrogen) and the Agilent 2100 Bioanalyzer with a DNA High Sensitivity Chip (Agilent) were used to assess the quality and concentration of libraries. The sequencing was performed by IMGM Laboratories GmbH (Martinsried) using a NovaSeq6000 sequencer (Illumina) equipped with NovaSeq6000 SP 100 v1.5 chemistry (Illumina). The goal was to get a total of 14 million readings per sample by using single-read sequencing with a read length of 100 base pairs. The raw sequencing data was subjected to processing and mapping using the Lexogen Bluebee pipeline. The differential expression analysis was performed using DESEQ2 algorithms on the Bluebee platform.

In the head, 113 genes were upregulated and 49 downregulated. In the abdominal fat body, 19 genes were upregulated and one gene downregulated. In the gut, we identified 55 upregulated genes and a single downregulated gene (Table S1).

### Histology and Imaging

#### Immunohistochemistry and confocal microscopy

To validate the expression of octopamine receptors in the fat body, we examined T2A-GAL4 insertions into receptor loci ^48^ using UAS-myr::RFP (BL# 7119). Adult flies were dissected in 4% PFA in PBS, and their head and abdomen were left in the fixative for 40 minutes at room temperature (RT), without agitation. Following fixation, these tissues were washed in PBST (0.1% Triton X-100 in PBS) for 10 minutes. The samples were then incubated in PBST with Hoechst 33258 solution (343-07961, Dojindo, 1:100) for 10 minutes to stain nuclei, followed by washing with PBST. For fluorescence imaging, SeeDB2-S ^75^ was used as the mounting medium. Images were acquired with the 40x/1.3 oil immersion objective (UPLFLN40XO, Olympus) using the Olympus FV1200 confocal microscope. Scan settings of the RFP channel were kept constant across specimens (Laser power: 8.5%; HV: 720) for intensity comparisons.

The immunohistochemistry methodology was derived from the work of Haiyang Chen, 2014^76^. The fat bodies of adult flies, along with the cuticles of their heads and abdomens, were dissected in PBS. The dissected tissues were fixed in 4% EM-grade paraformaldehyde buffer for 30 minutes. The buffer was composed of 100 mM glutamic acid, 25 mM KCl, 20 mM MgSO4, 4 mM Na2HPO4, 1 mM MgCl2, and had a pH of 7.4. Followed by washing three times, 15 minutes each, in a wash buffer (PBS with the addition of 0.3% Triton X-100), the tissues were blocked in the wash buffer containing 0.5% BSA for 15 minutes, and then exposed to primary antibodies diluted in the wash buffer at 4°C overnight. After receiving three washes in the wash buffer for 15 minutes each, the tissues were then exposed to secondary antibodies overnight at room temperature. Next followed by repeating the aforementioned washing procedures three times for 15 minutes each, and then performing an additional one-hour wash. Finally, the mounting method was carried out. The primary antibodies employed in the study were rabbit anti-GFP (Invitrogen A11122, 1:200), mouse anti-RFP (Takara 632393, 1:200), rat anti-NC82 (DSHB 528121, 1:200), and mouse anti-GFP (Neuromab, c/o antibodies inc. 75-132, 1:200). The second antibodies employed were goat anti-rabbit 488 (Invitrogen A11034, 1:200), goat anti-mouse 594 (Invitrogen A11005, 1:200), goat anti-rat 647 (Invitrogen A48265, 1:200), and goat-anti-mouse 488 (Invitrogen A11001, 1:500). The nucleus was labeled with TOPRO3 (Invitrogen T3605, 1:200).

All immunofluorescence images were acquired using a Leica SP5 confocal microscope. Images were collected for each set of trials using identical settings. The image data was processed using ImageJ.

### CaMPARI2 experiments

#### OA-stimulation tests

Seven-day-old mated female flies with the genotype *Cg-Gal4;UAS-CaMPARI2* were collected, and the fat bodies attached to the cuticles from the abdomen and head were then dissected in a saline solution. The fat body samples were thereafter subjected to either 5 µg/mL octopamine in saline (test group) or saline alone (control group) for a period of 8 to 10 minutes at room temperature. After that, a 405 nm ultraviolet (UV) light was applied to the tissue from a distance of 3 cm for a period of 30 seconds using a 405 nm light-emitting diode (LED) stand light with an intensity of 200 mW/cm² (M395L5 model by Thorlabs). Following UV stimulation, images of the fresh fat bodies were promptly captured using a Leica SP5 confocal microscope equipped with a 20x water objective. To enhance the fluorescence conversion of CaMPARI2, we employed a 405 nm laser from the Leica SP5 along with a 10% intensity 488 nm laser and a 15% intensity 561 nm laser. To provide fair comparisons and minimize potential biases, fresh octopamine solutions were prepared for each experimental day, and imaging was performed a maximum of two times on the same tissue. The imaging configuration remained identical.

#### Optogenetic Tdc2 neural stimulation tests

Freshly hatched flies with the genotype *Cg-Gal4/UAS-CaMPARI2;Tdc2-lexA/LexAop-CsChrimson* were collected and kept on retinal food until they were 7 days old. Mated female flies were then collected in darkness. For the test group, each awake fly was placed in a 200 µL pipette tip and exposed to a 660 nm LED stand light (3 mW/mm², M660F1, Thorlabs) from 1 cm above for 3 minutes to activate the Tdc2 neurons. Immediately after the light exposure, the flies were anesthetized on ice, and the fat bodies from the abdomen and head were dissected in saline for further light exposure treatment. The test group flies were subjected to alternating cycles of red light and UV light (200 mW/cm², M395L5, Thorlabs), consisting of three cycles of 15 seconds with red light on and 15 seconds off, while the UV light remained constantly on for 90 seconds. In the ‘no red light’ control group, no red light was applied before dissection nor after; instead, only 405 nm light irradiation was administered on the fat body tissue for 90 seconds as a light control. Following light stimulation and/or photoconversion, the fat bodies were promptly moved to a glass slide and then covered with a cover slip. Images of the fresh fat bodies were then acquired using a Leica SP5 confocal microscope with a 20x water objective. In order to optimize the fluorescence conversion of CaMPARI2, a laser with a wavelength of 405 nm at an intensity of 5% was used in conjunction with a laser with a wavelength of 488 nm at an intensity of 10% and a laser with a wavelength of 561 nm at an intensity of 15%. To assure unbiased comparisons, 5 µg/mL octopamine solutions were made freshly on each experimental day, and imaging was conducted a maximum of two times on the same tissue. The microscope parameter settings were consistent across all experimental groups.

Maximum intensity projections were generated from Z-stacks using ImageJ. CaMPARI2 photoconversion rates were quantified by calculating the Red/Green ratio in ImageJ and Excel. Regions of interest (ROIs) were drawn around fat body cells in the green channel and copied to the background to measure the average green intensity. The same ROIs were applied to the red channel to obtain background-subtracted red intensity. The average green and red signal intensities from the cell bodies and background were computed, and the Red/Green ratio was calculated using the following formula:

Red/Green ratio = (Average red signal intensity of cell bodies – Average red signal intensity of the background) / (Average green signal intensity of cell bodies – Average green signal intensity of the background). Each representative image was cropped around the cell bodies.

#### Grab-DA experiments

Seven-day-old mated female flies with the genotype Cg-Gal4;UAS-GrabDA, UAS-myr-mRFP were collected, and fat bodies attached to the abdominal cuticles were dissected in saline solution. The dissected fat bodies were mounted on slides containing either 5 µg/mL octopamine in saline (test group) or saline alone (control group). Fresh tissue images were immediately captured using a Leica SP5 confocal microscope equipped with a 20x water objective. To ensure consistency and minimize bias, fresh octopamine solutions were prepared daily, and each tissue was imaged only once under identical imaging settings.

Maximum intensity projections were generated from Z-stacks using ImageJ, and Grab-DA fluorescent readout quantification was performed by calculating the Green/Red fluorescence ratio. The average intensities of green and red signals were measured from the cell bodies and background using ImageJ and Excel. The Green/Red ratio was determined using the formula: Green/Red ratio = (Average green signal intensity of cell bodies – Average green signal intensity of the background) / (Average red signal intensity of cell bodies – Average red signal intensity of the background). Each representative image was cropped to focus on the cell bodies.

### Data collection and statistical analyses

The statistical analyses were conducted using the GraphPad Prism 10 software. The normality of the datasets was assessed using the D’Agostino & Pearson, Anderson-Darling, Shapiro-Wilk, Kolmogorov-Smirnov tests, and QQ plot. The Wilcoxon matched-pairs signed rank test was employed to examine the total number of sips of the two food substrates in the flyPAD and optoPAD trials. The RNA-seq data was analyzed using the DESeq2 algorithm on the Bluebee platform. A cut-off of 0.1 was applied to the adjusted p-value in order to identify genes that showed substantial upregulation or downregulation. The CaMPARI2 data were evaluated by calculating the ratio of red to green CaMPARI signals using the Mann-Whitney test. The Grab-DA data were examined by calculating the ratio of green Grab-DA signals to red myr::RFP in the fat body using the Welch-corrected t-test.

The significance level for all the data, excluding RNA-seq data, was established at 0.05, in accordance with recognized statistical conventions. The relevant notations are: ns (p > 0.05), * (p < 0.05), ** (p < 0.01), *** (p < 0.001), and **** (p < 0.0001). The error bars in the graphs represent the standard error of the mean (SEM). Boxplots employ the Tukey method to calculate the length of the whiskers, which is equivalent to 1.5 times the interquartile range (IQR). Data points that reside outside this range are classified as outliers and are shown separately.

The sample size n in all experiments denotes the number of flies or tissue samples. Each experiment was carried out at least twice independently with different parental animals on different days to ensure repeatability. For all experiments, controls and test groups were run side-by-side and the order and position of the behavioral arenas used in the flyPAD was randomized. All data were used for statistical analysis.

