## Supplementary figures for "A Bidirectional Brain-Fat Body Axis for Pathogen Avoidance"

Figures S1 to S4
Table S1

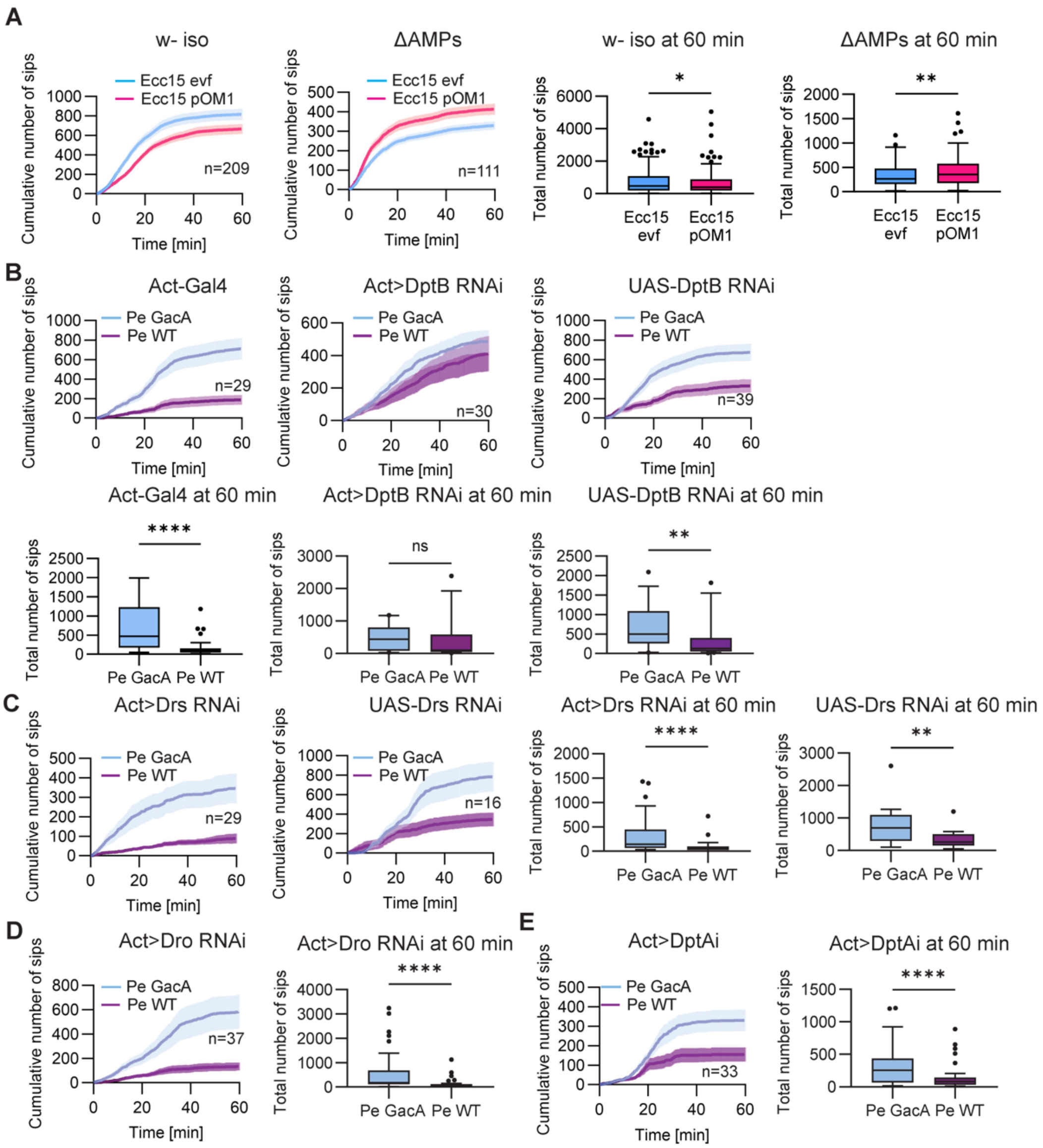

A
w- iso
ΔAMPs
w- iso at 60 min
ΔAMPs at 60 min
Ecc15 evf
Ecc15 pOM1
n=209
n=111
Cumulative number of sips
Total number of sips
Time [min]
B
Act-Gal4
Act>DptB RNAi
UAS-DptB RNAi
Act-Gal4 at 60 min
Act>DptB RNAi at 60 min
UAS-DptB RNAi at 60 min
Pe GacA
Pe WT
n=29
n=30
n=39
ns
C
Act>Drs RNAi
UAS-Drs RNAi
Act>Drs RNAi at 60 min
UAS-Drs RNAi at 60 min
n=29
n=16
D
Act>Dro RNAi
Act>Dro RNAi at 60 min
n=37
E
Act>DptAi
Act>DptAi at 60 min
n=33

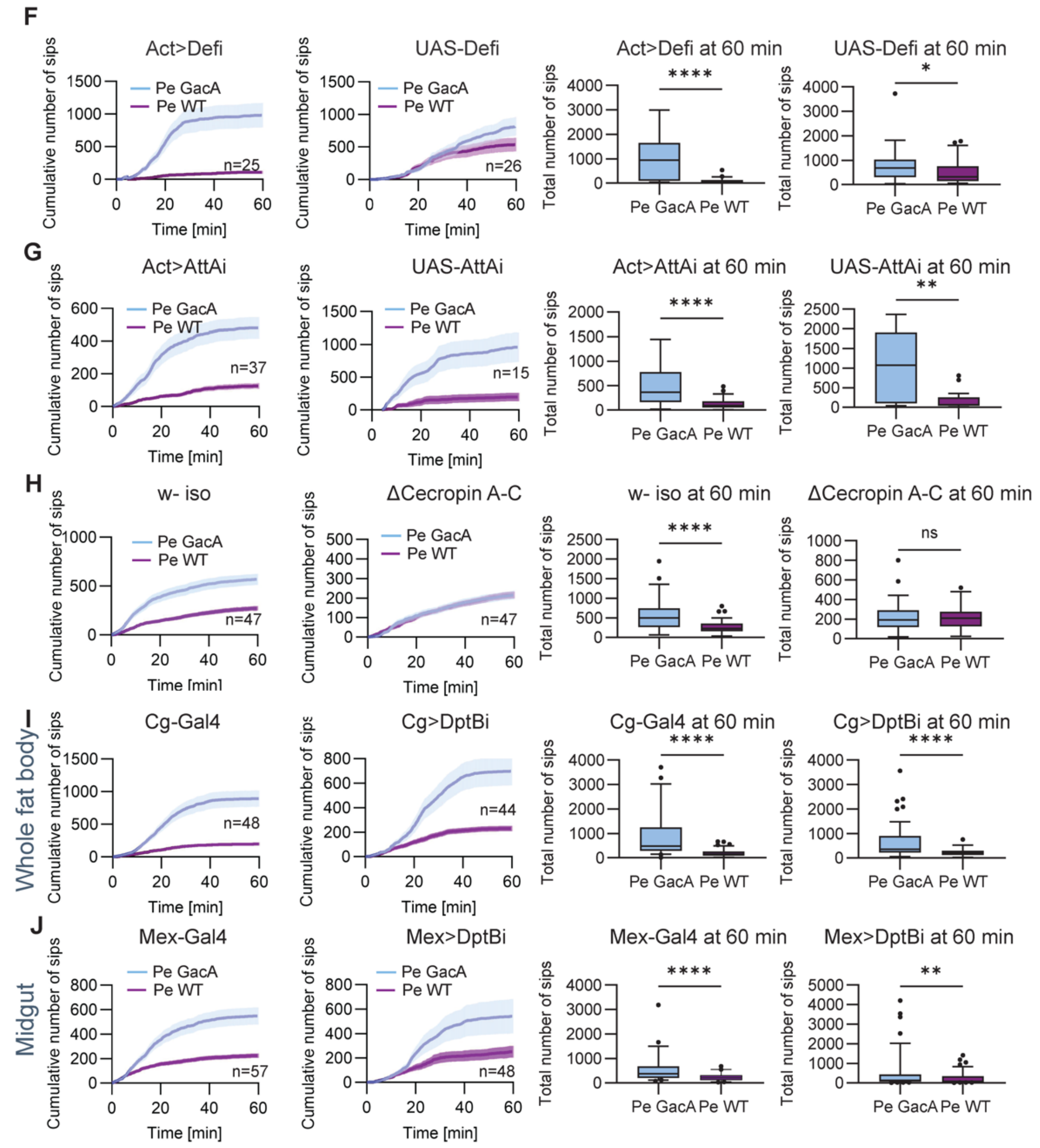


**Figure S1**

(A) Feeding preferences of starved, naïve isogenic *w-* flies and ΔAMPs flies for the harmless and harmful *Ecc15* bacteria: n = 209 (control w-), n = 111 (ΔAMPs). Cumulative number of sips; mean ± SEM. Total number of sips at 60 mins; *p-values* calculated via Wilcoxon matched-pairs signed rank test.

(B) Feeding preferences for harmless and harmful *Pe* bacteria upon global knock-down of DptB using Actin-Gal4;UAS-DptB RNAi (n = 30), and the control flies Actin-Gal4 (n = 29), UAS-DptB RNAi (n = 39); experiments at 28°C. Cumulative number of sips; mean ± SEM. Total number of sips at 60 mins; *p-values* calculated via Wilcoxon matched-pairs signed rank test.

(C) Feeding preferences for harmless and harmful *Pe* bacteria upon global knock-down of Drosomycin using Actin-Gal4;UAS-Drs RNAi (n = 29), and the control flies UAS-Drs RNAi (n = 16); experiments at 28°C. Cumulative number of sips; mean ± SEM. Total number of sips at 60 mins; *p-values* calculated via Wilcoxon matched-pairs signed rank test.

(D) Feeding preferences for harmless and harmful *Pe* bacteria upon global knock-down of Drosocin using Actin-Gal4;UAS-Dro RNAi (n = 37); experiments at 28°C. Cumulative number of sips; mean ± SEM. Total number of sips at 60 mins; *p-values* calculated via Wilcoxon matched-pairs signed rank test.

(E) Feeding preferences for harmless and harmful *Pe* bacteria upon global knock-down of Diptericin A using Actin-Gal4;UAS-DptA RNAi (n = 33); experiments at 28°C. Cumulative number of sips; mean ± SEM. Total number of sips at 60 mins; *p-values* calculated via Wilcoxon matched-pairs signed rank test.

(F) Feeding preferences for harmless and harmful *Pe* bacteria upon global knock-down of Defensin using Actin-Gal4;UAS-Def RNAi (n = 25) and control flies UAS-Def RNAi (n = 26); experiments at 28°C. Cumulative number of sips; mean ± SEM. Total number of sips at 60 mins; *p-values* calculated via Wilcoxon matched-pairs signed rank test.

(G) Feeding preferences for harmless and harmful *Pe* bacteria upon global knock-down of Attacin A using Actin-Gal4;UAS-AttA RNAi (n = 37) and control flies UAS-AttA RNAi (n = 15); experiments at 28°C. Cumulative number of sips; mean ± SEM. Total number of sips at 60 mins; *p-values* calculated via Wilcoxon matched-pairs signed rank test.

(H) Feeding preferences for harmless and harmful *Pe* bacteria for Cecropins mutant flies (ΔCecropin A-C) and control flies: n = 47 (Δ Cecropin A-C), n = 47 (isogenic *w-*). Cumulative number of sips; mean ± SEM. Total number of sips at 60 mins; *p-values* calculated via Wilcoxon matched-pairs signed rank test.

(I) Feeding preferences for harmless and harmful *Pe* bacteria upon knock-down of DptB in fat body using Cg-Gal4;UAS-DptB RNAi (n = 44) and control flies Cg-Gal4 (n = 48). Cumulative number of sips; mean ± SEM. Total number of sips at 60 mins; *p-values* calculated via Wilcoxon matched-pairs signed rank test.

(J) Feeding preferences for harmless and harmful *Pe* bacteria upon knock-down of DptB in epithelial cells in the midgut using Mex-Gal4;UAS-DptB RNAi (n = 48) and control flies Mex-Gal4 (n = 57), experiments at 28°C. Cumulative number of sips; mean ± SEM. Total number of sips at 60 mins; *p-values* calculated via Wilcoxon matched-pairs signed rank test.

For all analyses, the statistical notation is as follows: ns (not significant), $p > 0.05$; * $p < 0.05$; ** $p < 0.01$; *** $p < 0.001$; **** $p < 0.0001$. Error bars in all panels represent the standard error of the mean (SEM).

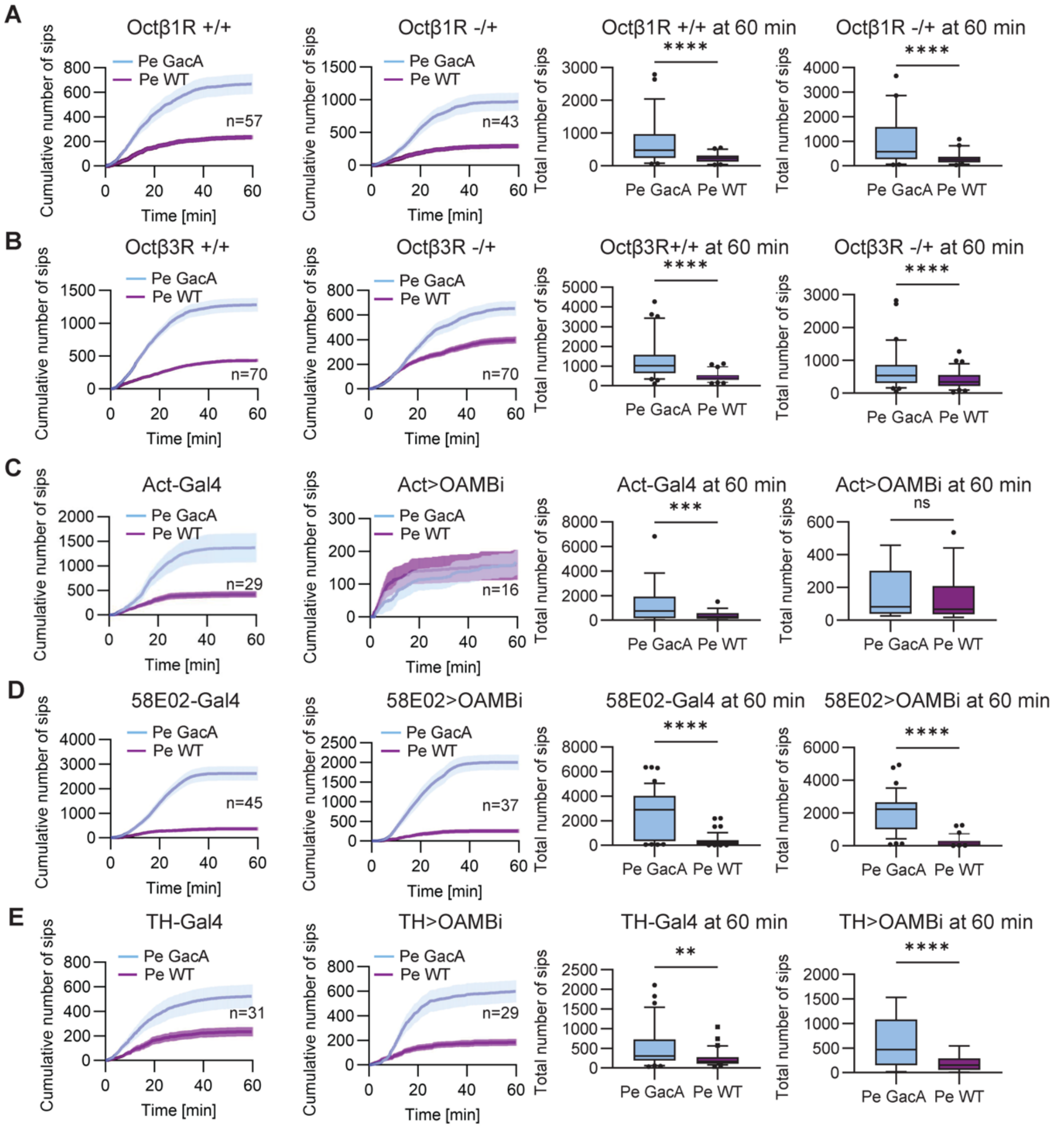


**Figure S2**

(A) Feeding preferences for harmless and harmful *Pe* bacteria for heterozygous Octβ1R mutant flies (Octβ1R-/+) and control flies: n = 43 (Octβ1R-/+), n = 57 (Octβ1R+/+). Octβ1R homozygous is lethal. Cumulative number of sips; mean ± SEM. Total number of sips at 60 mins; *p-values* calculated via Wilcoxon matched-pairs signed rank test.

(B) Feeding preferences for harmless and harmful *Pe* bacteria for heterozygous Octβ3R mutant flies (Octβ3R-/+) and control flies: n = 70 (Octβ3R-/+), n = 70 (Octβ3R+/+). Octβ3R homozygosity is lethal. Cumulative number of sips; mean ± SEM. Total number of sips at 60 mins; *p-values* calculated via Wilcoxon matched-pairs signed rank test.

(C) Feeding preferences for harmless and harmful *Pe* bacteria upon global knock-down of OAMB using Actin-Gal4;UAS-OAMB RNAi (n = 16), and the control flies Act-Gal4 (n = 29). Cumulative number of sips; mean ± SEM. Total number of sips at 60 mins; *p-values* calculated via Wilcoxon matched-pairs signed rank test.

(D) Feeding preferences for harmless and harmful *Pe* bacteria upon knock-down of OAMB in PAM cluster neurons using 58E02-Gal4;UAS OAMB RNAi (n = 37), and the control flies 58E02-Gal4 (n = 45). Cumulative number of sips; mean ± SEM. Total number of sips at 60 mins; *p-values* calculated via Wilcoxon matched-pairs signed rank test.

(E) Feeding preferences for harmless and harmful *Pe* bacteria upon knock-down of OAMB in PPL1 cluster neurons using TH-Gal4;UAS-OAMB RNAi (n = 29), and the control flies TH-Gal4 (n = 31). Cumulative number of sips; mean ± SEM. Total number of sips at 60 mins; *p-values* calculated via Wilcoxon matched-pairs signed rank test.

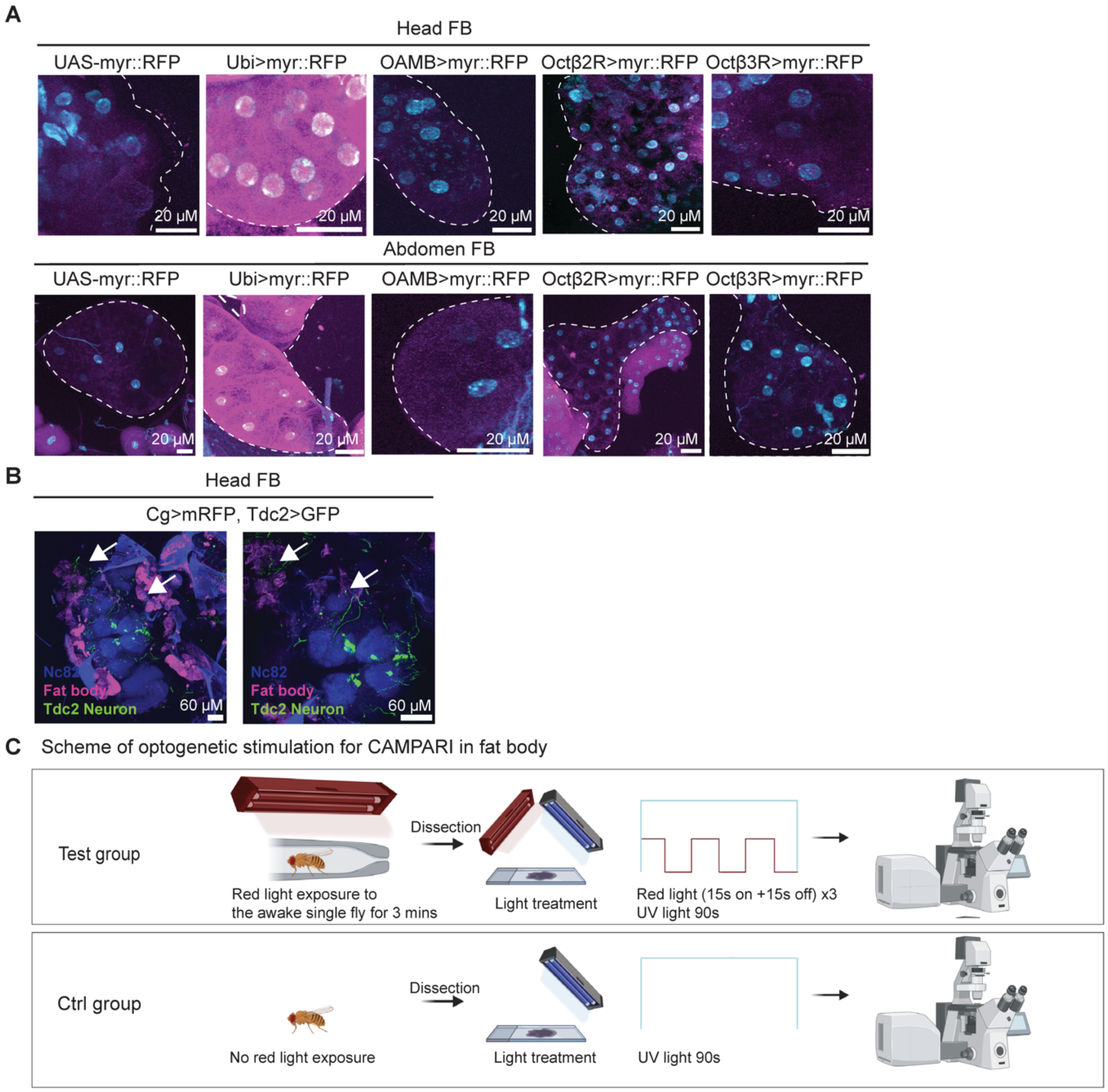


**Figure S3**

(A) Representative Z-projection images of fat bodies of three octopamine receptor reporter knock-in lines: OAMB-T2A-Gal4;UAS-myr::mRFP, Octβ2R-T2A-Gal4;UAS-myr::mRFP, and Octβ3R-T2A-Gal4;UAS-myr::mRFP with respective negative controls (UAS-myr::mRFP) and positive controls (Ubi-Gal4;UAS-myr::RFP). Nuclei are labeled with Hoechst33342 (cyan). Scale bar: 20 μm.

(B) Example Z-projection images of head fat body and OANs in flies (Cg-Gal4, UAS-RFP; Tdc2-lexA, lexAop-GFP) showing anatomical proximity of OANs (green) with head fat body (magenta). Nc82 was used as counterstaining for the fly brain (blue). OANs are located adjacent to the foramen where the esophagus traverses the brain. Scale bar = 20 μm.

(C) Schematic of optogenetic stimulation for CaMPARI2 experiments.

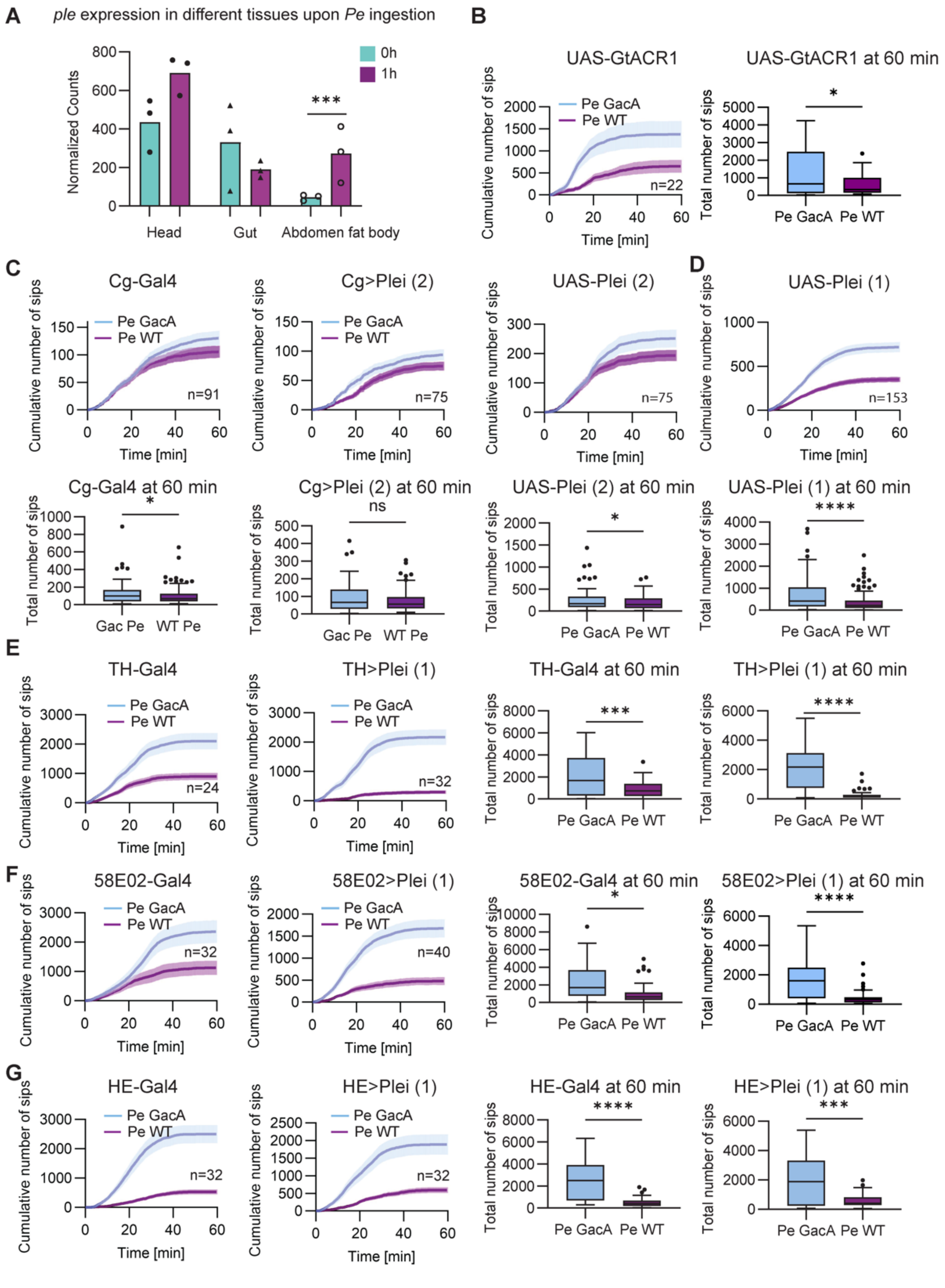


**Figure S4**

(A) Boxplots depicting normalized counts from DESeq2 for *ple* expression level before and after ingesting *Pe WT* for 1 hour. *Ple* was significantly upregulated in the abdominal fat body upon *Pe WT* ingestion. The adjusted p-values were obtained from DESeq2 analysis.

(B) Feeding preferences for harmless and harmful *Pe* bacteria for optogenetic control flies UAS-GtACR1 (n = 22). Cumulative number of sips; mean ± SEM. Total number of sips at 60 mins; *p-value* calculated via Wilcoxon matched-pairs signed rank test.

(C) Feeding preferences for harmless and harmful *Pe* bacteria upon knock-down tyrosine hydroxylase in fat body using another UAS-Ple RNAi line (BDSC# 65875) Cg-Gal4;UAS-Ple RNAi (n = 75), and control flies Cg-Gal4 (n = 91), UAS-ple RNAi (n = 75). Cumulative number of sips; mean ± SEM. Total number of sips at 60 mins; *p-values* calculated via Wilcoxon matched-pairs signed rank test.

(D) Feeding preferences for harmless and harmful *Pe* bacteria for control flies UAS-Ple RNAi (BDSC# 76069) (n = 153). Cumulative number of sips; mean ± SEM. Total number of sips at 60 mins; *p-values* calculated via Wilcoxon matched-pairs signed rank test.

(E) Feeding preferences for harmless and harmful *Pe* bacteria upon knock-down tyrosine hydroxylase in PPL1 neurons using TH-Gal4;UAS-Ple RNAi (1) (n = 32), and control flies TH-Gal4 (n = 24). Cumulative number of sips; mean ± SEM. Total number of sips at 60 mins; *p-values* calculated via Wilcoxon matched-pairs signed rank test.

(F) Feeding preferences for harmless and harmful *Pe* bacteria upon knock-down tyrosine hydroxylase in PAM neurons using 58E02-Gal4;UAS-Ple RNAi (1) (n = 40), and control flies 58E02-Gal4 (n = 32). Cumulative number of sips; mean ± SEM. Total number of sips at 60 mins; *p-values* calculated via Wilcoxon matched-pairs signed rank test.

(G) Feeding preferences for harmless and harmful *Pe* bacteria upon knock-down tyrosine hydroxylase in hemocytes using HE-Gal4;UAS-Ple RNAi (1) (n = 32), and control flies HE-Gal4 (n = 32). Cumulative number of sips; mean ± SEM. Total number of sips at 60 mins; *p-values* calculated via Wilcoxon matched-pairs signed rank test.

For all analyses, the statistical notation is as follows: ns (not significant), $p > 0.05$; * $p < 0.05$; ** $p < 0.01$; *** $p < 0.001$; **** $p < 0.0001$. Error bars in all panels represent the standard error of the mean (SEM).
